# Cross-expression meta-analysis of 695 brain samples reveals coordinated gene expression across spatially adjacent cells

**DOI:** 10.1101/2024.09.17.613579

**Authors:** Ameer Sarwar, Mara Rue, Leon French, Helen Cross, Sarah Choi, Xiaoyin Chen, Jesse Gillis

## Abstract

Spatial transcriptomics promises to transform our understanding of tissue biology by molecularly profiling individual cells *in situ*. A fundamental question they allow us to ask is how nearby cells orchestrate their gene expression. Rather than focus on how these cells (samples) communicate with each other, we reframe the problem to investigate how genes (features) coordinate their expression between neighboring cells. To study these phenomena – called cross-expression – we compare all genes to find pairs that coordinate their expression between adjacent cells, thereby avoiding curating gene lists or annotating cell types. Our end-to-end method recovers ligand-receptor pairs as cross-expressing genes and finds gene combinations that mark anatomical regions, complementing marker gene-based region annotation. Leveraging the overlapping genes across different panels, we use multiple atlas-scale adult mouse brain datasets (~25 million cells, 695 samples, 8 technologies) to create an integrated, meta-analytic cross-expression network, whose communities are enriched in spatial processes such as synaptic signaling and G protein coupled receptor activity. Highlighting cross-expression’s biological utility, our network shows that genes *Drd1* and *Gpr6*, which are individually implicated in Parkinson’s disease (PD) and are being pursued as therapeutic targets, are cross-expressed within the striatum, hinting at their joint role in PD pathophysiology. We provide an efficient R package (https://github.com/gillislab/CrossExpression/) to computationally analyze and visually explore cross-expression patterns, which allow us to better understand how genes coordinate their expression in space to perform tissue-level functions.

## Background

Spatial transcriptomic technologies record cells’ gene expression alongside their physical locations, enabling us to understand how they influence one another within the tissue [1]. Focusing on select genes, imaging-based platforms profile expression at the single cell level, giving us a high-resolution snapshot of spatial gene expression [2–8]. They have facilitated numerous studies on defining local spatial patterns [9–12], finding gene covariation in spatial niches [13–19], elucidating cell-cell interactions using ligand-receptor expression [20–28], and determining spatial cell type heterogeneity and tissue structure [29–31]. These efforts have resulted in a greater understanding of tissue biology, culminating in the generation and exploration of reference atlases [32–35].

Imaging-based platforms can now profile over a thousand genes in millions of cells [2–8], generating large amounts of data ripe for biological discovery. These data can be analyzed at a high resolution, where we can investigate individual genes, single cells, and their spatial relations. The dominant framework in this space is the cell-cell communication methods [20–28], which infer co-localized ligands and receptors in neighboring cells. These approaches typically rely on curated ligand-receptor databases to infer interactions between cell types. Despite rapid progress in this area, these methods have important limitations. First, the reliance on extant databases limits novel discoveries to well-known ligands and receptors, thereby overlooking potential interaction partners present in the gene panel. Second, the use of cell types requires that the annotations are sufficiently accurate, but this is currently an open problem owing to the limited size of the gene panels, often requiring ‘label transfer’ from matched single-cell RNA-seq datasets. Indeed, the communication inference is made at the aggregate, cell type level when the underlying data allows a fully ‘bottom-up’ analysis at the single cell level. Third, the cell-centric perspective makes it difficult to integrate datasets across experiments, limiting the inferences to single studies. Because large-scale datasets are rapidly being collected, an approach is needed to develop a ‘common map’ between them to discover reproducible biological signals.

The gene-centric framework is an alternative to the cell-cell inference approaches. By prioritizing genes (features) over cells (samples), it allows comparisons across datasets using genes shared between their panels. Moreover, when the cell-cell interaction methods use ligand-receptor co-localization, they implicitly leverage the gene-centric perspective, where inferences are made if the relevant genes are expressed in neighboring cells more frequently than expected by chance. A number of studies have explicitly used the gene-centric approach. For example, Haviv et al. (2024) model how gene-gene co-expression within the same cells changes across spatial niches [14]. Studies by Miller et al. (2021) and Li et al. (2023) consider gene-gene coordination between spatially adjacent cells using the global bivariate Moran’s *I* statistic [18,19]. Whereas Haviv et al. (2024) ignore gene-gene coordination between neighboring cells, Miller et al. (2021) and Li et al. (2023) require ligand-receptor databases, thus limiting their scope. Additionally, the spatial correlation methods, such as the global bivariate Moran’s *I* statistic, are known to increase false positives because they do not account for “within location” association. For example, if genes A and B are both co-expressed in neighboring cells, then they would appear correlated across neighbors even when the two cells function fully autonomously.

Here, we introduce “cross-expression”, which models the degree to which genes coordinate their expression across neighboring cells. Avoiding the use of curated databases and cell type labels, our fully end-to-end and bottom-up method uses the raw gene expression and cell location matrices to determine which gene pairs, among all possible pairs, coordinate their expression between neighboring cells. We explicitly account for spurious associations induced by “within cell” co-expression patterns, thereby revealing genuine coordination once cell intrinsic processes have been controlled. A consequence of our method is that it recovers ligand-receptor pairs as cross-expressed genes, e.g., *Sst* and *Sstr2*, thus extending the well-developed cell-cell communication framework. Its gene-centric approach facilitates integrative analysis, where we create a cross-expression meta-analytic network using 13 datasets spanning ~25 million cells, 695 brain samples, and 8 technological platforms. Our network shows highly modular structure, with communities enriched for gene ontology (GO) terms like synaptic signaling, neurotransmitter regulation, cell adhesions, G protein receptor activity, and other spatial processes. Highlighting its biological utility, the network reveals that across numerous samples the genes *Drd1* and *Gpr6*, which are individually known to play roles in Parkinson’s disease (PD) and are being pursued as therapeutic targets, are cross-expressed in the striatum, hinting at their joint function in the central anatomical locus in PD pathophysiology. We also show that the cross-expression patterns in one dataset reliably predict those in other datasets as well as that similar anatomical regions have similar cross-expression profiles, indicating that our method reliably detects subtle variations in gene expression across batches and anatomical space. To facilitate the analysis and exploration of cross-expression patterns, we provide a fully vectorized R package that processes hundreds of thousands of gene pairs across millions of cells within minutes on a standard personal laptop, allowing in-depth analyses of how genes coordinate their expression in space to perform tissue-level functions.

## Results

### Cross-expression overview

Cross-expression is defined as the degree to which two genes coordinate their expression across neighboring cells (Fig. 1a). Specifically, it asks whether the expression of gene A in a cell is associated with the expression of gene B in the cell’s (nearest) neighbor. To properly capture these patterns, we control for co-expression or, more generally, the association induced between neighboring locations due to the association present within the same location. In particular, if genes A and B are co-expressed in both the neighboring cells, then they would appear correlated across these cells even when the cells function completely independently. We control for this by defining cross-expression mutually exclusively, namely gene A is expressed in the cell without gene B and gene B is expressed in the neighbor without gene A.

**Fig. 1:**
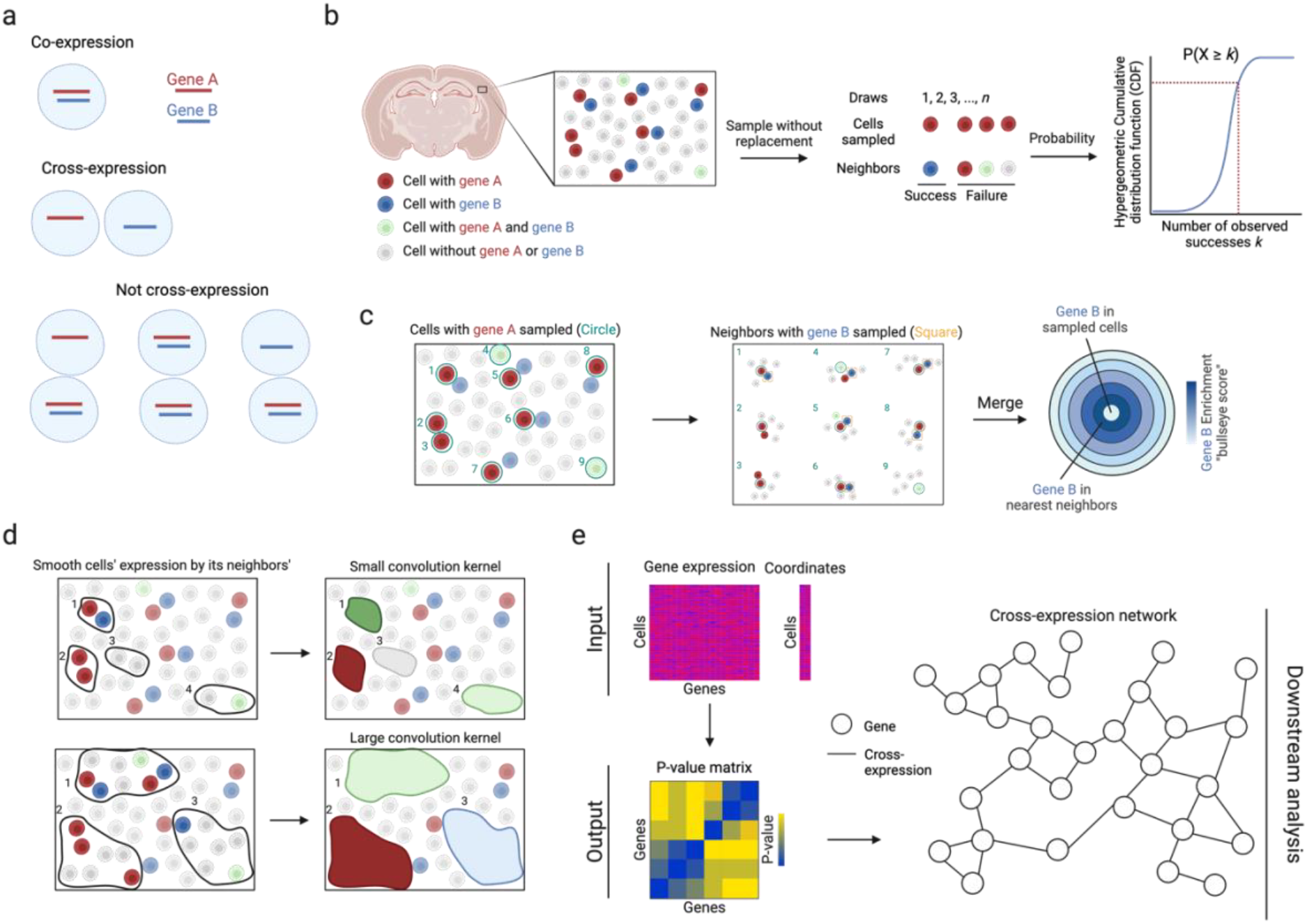
Cross-expression analysis. **a**, Cross-expression is the mutually exclusive expression of genes between neighboring cells. If either cell expresses both genes, the cell pair is not considered to cross-express. **b**, The probability that two genes cross-express is modeled by the hypergeometric distribution, where the ‘successful trials’ are the cell-neighbor pairs where the cell expresses gene A and the neighbor expresses gene B. **c**, Cross-expression is compared to co-expression to quantify the effect size, where the number of neighbors with gene B is compared to the number of cells co-expressing genes A and B. ‘Sampled cells’ (center) are those expressing gene A and neighbors are concentric rings, with the order indicating the *n*-th neighbor. **d**, Averaging gene expression between cells and their neighbors smooths it, extending cross-expression analysis from cell pairs to regions. Number of neighbors is the kernel size. **e**, Software inputs are the gene expression and cell location matrices, and the output is a p-value matrix, which enables downstream analyses, such as the creation of cross-expression networks. Created with BioRender.com.

We quantify cross-expression in three statistically robust and highly interpretable ways. First, we use the hypergeometric test to compute p-values (Fig. 1b). We begin by finding each cell’s nearest neighbor to create a cell-to-neighbor ‘mapping’. While cells almost never have two nearest (equidistant) neighbors, two or more cells can have the same neighbor. The hypergeometric test is performed on these cell-neighbor pairs. For genes A and B, we count the number of pairs where the cell expresses gene A (sample size) and the number of pairs where the neighbor expresses gene B (possible successes). We then compute the intersection (observed successes), namely the number of pairs where the cell expresses A and the neighbor expresses B. Together with the total number of pairs (population size), we use the hypergeometric cumulative distribution function (CDF) to analytically compute the probability of observing as many or more successes, giving us a p-value for the cross-expression of genes A and B.

Second, we quantify the effect size by comparing the number of cross-expressing pairs to the number of co-expressing cells (Fig. 1c). The central idea is that if gene B is randomly assigned to cells while gene A is held constant, then we expect some co-expression and some cross-expression simply by chance, where their expected ratio is 1. This ratio is our effect size, with values greater than 1 indicating more cross-than co-expression, and we represent it till the *n*-th neighbor using concentric rings in the bullseye plot.

Third, we compute a Pearson correlation between gene A in each A-expressing cell and gene B in its nearest neighbor. This yields a directed set of cell-neighbor pairs, capturing how B varies across the local context of A. Some neighbors are shared by multiple cells, reflecting spatial centrality; this weighting mirrors the biological structure of the tissue, where certain cells may influence more of their surroundings. Although this induces non-uniform contributions, each pair is treated independently from the target cell’s perspective, preserving statistical validity for correlation.

Although we focus on individual cells, groups of cells may form spatial niches and gene expression may be coordinated between niches. To assess cross-expression at this coarser resolution, we average a gene’s expression in a cell with its expression in the neighbors (Fig. 1d), thus smoothing it within a spatial niche, with the number of neighbors forming the niche size. Accordingly, cross-expression can be compared across niches by, for example, finding associations between smoothed niche-specific gene expression profiles.

To enable these analyses, we provide an efficient software package in R that requires the gene expression and cell location matrices as inputs, and outputs a gene-gene p-value matrix that facilitates downstream analyses, such as the creation of cross-expression networks (Fig. 1e). The package also contains functions for computing effect sizes, making bullseye plots, smoothing gene expression, viewing cross-expressing cells *in situ*, and assessing if cross-expression is spatially enriched. Collectively, the cross-expression framework uses spatial information to discover how genes coordinate their expression across neighboring cells, thereby providing a useful analytical framework for deeply exploring spatial transcriptomic data.

### Cross-expression recovers ligand-receptor pairs and reveals coordinated gene expression profiles across the tissue

To study cross-expression, we used BARseq (barcoded anatomy resolved by sequencing) to collect data from a whole mouse brain. This dataset profiled expression in 1 million cells across 16 sagittal slices, using a gene panel of 104 cortical cell type markers and 25 ligands and receptors, including neuropeptides, their receptors, and monoamine neuromodulatory receptors. Because receptors and the enzymes that synthesize the corresponding ligands are often expressed in nearby cells [20–28], we reasoned that these genes should show cross-expression. As an example, we find that across the cortical somatosensory nose region and visceral areas (Fig. 2a), the neuropeptide somatostatin *Sst* and its cognate receptor *Sstr2* are cross-expressed (Fig. 2b, left, p-values ≤ 0.01 and 0.05, respectively). Indeed, these genes are consistently expressed across neighboring cells (Fig. 2b, right), a pattern that is otherwise difficult to discover without prior knowledge. While cross-expression recovers ligand-receptor genes, it does not imply that they form synaptic or endocrine connections, since spatial transcriptomic measurements are restricted to the cell body.

**Fig. 2:**
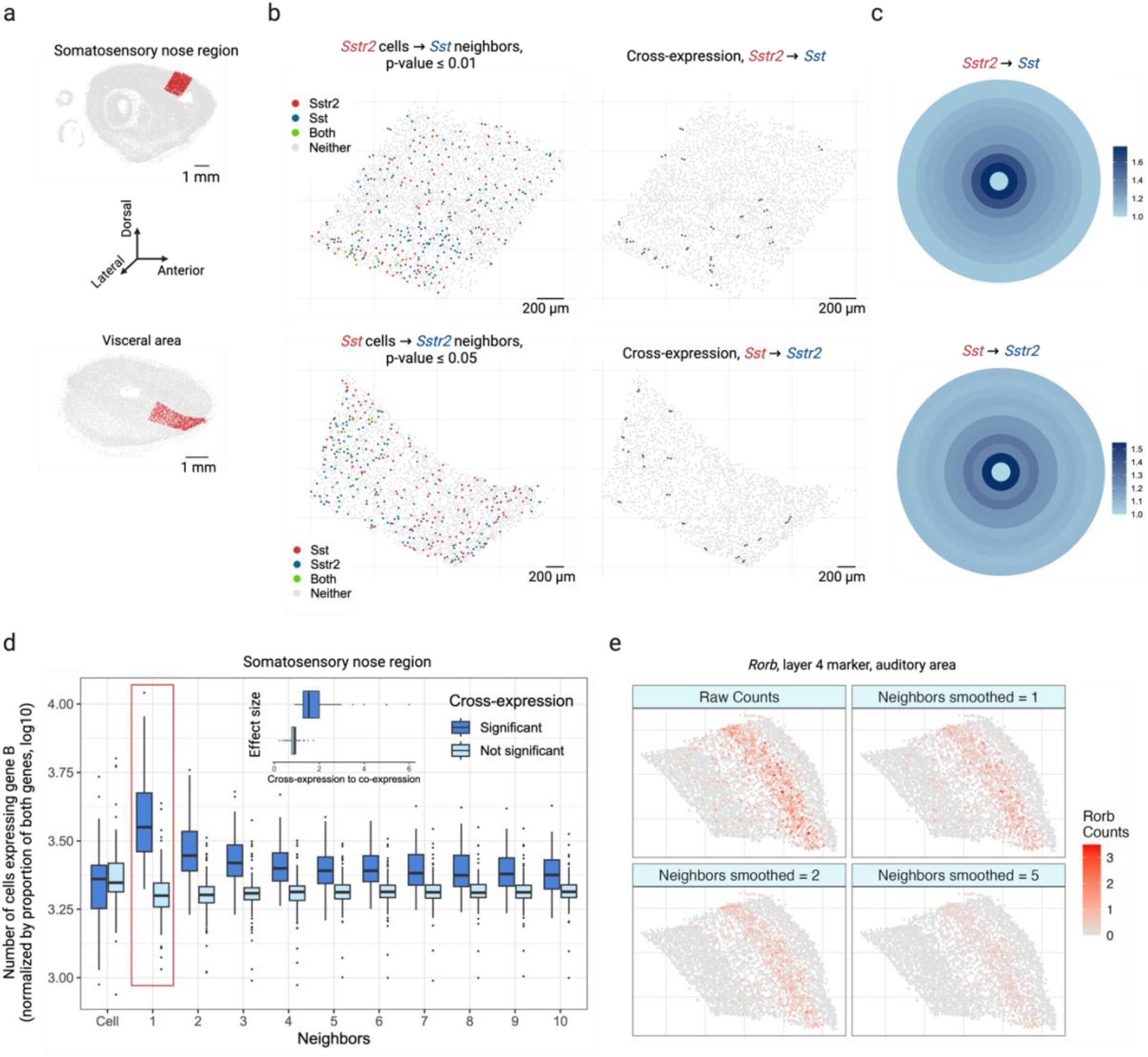
Cross-expression analysis reveals coordinated gene expression between neighboring cells. **a**, Sagittal brain slices showing cortical somatosensory nose region (top) and visceral area (bottom) as randomly selected regions of interest. **b**, Neuropeptide somatostatin *Sst* and its cognate receptor *Sstr2* cross-express in regions shown in (a). Points indicate cells and colors indicate gene expression (left), with cross-expressing cell pairs highlighted (right). **c**, Bullseye scores for *Sst* and *Sstr2* in the regions shown in (a, b). The scores are reported as ratio of cross-to co-expression. **d**, Bullseye scores for cross-expressing (significant) and non-cross-expressing (not significant) gene pairs in the somatosensory nose region. ‘Cell’ corresponds to the central ring in (c), and the red rectangle highlights the first neighbor/ ring. Inset, ratio of bullseye scores for the first neighbor to the central cells for cross-expressing and non-cross-expressing genes. Central line, median; box limits, first and third quartiles; whiskers, ±1.5x interquartile range; points, outliers. **e**, Smoothed gene expression for different numbers of neighbors for the auditory cortical layer 4 marker gene *Rorb*. Created with BioRender.com.

Next, we explore the bullseye plots, which allow us to quantify the effect size by comparing cross-expression with co-expression. For *Sst* and *Sstr2* in the somatosensory nose (2,015 cells) and visceral regions (1,603 cells), we see a bullseye pattern with low co-expression and high cross-expression that decreases for distant neighbors (Fig. 2c). Specifically, for these regions the bullseye score ratio between the first neighbor and the central cell is 1.8 and 1.6, respectively, whereas the ratio between the averaged second-to-tenth neighbor and the central cell is 1.3 and 1.2. These findings suggest that for central cells expressing one gene in a pair, a higher proportion of adjacent neighbors, but not the more distant ones, express the cognate gene within the local spatial niche, underscoring the specificity and resolution with which patterns of coordinated gene expression can be recovered. We next compare the bullseye plots for gene pairs with and without cross-expression (Fig. 2d), finding that the former match the patterns just described. To quantify this, we compare the bullseye scores of the nearest neighbors with those of cells expressing gene A, discovering that this ratio is much greater for genes that cross-express than for those that do not (Fig. 2d, inset, Mann-Whitney U test, p-value ≤ 0.001, median ratios: 1.5 and 0.9, respectively). Notably, this ratio is approximately 1 for genes that do not cross-express, suggesting that here gene B is expressed in neighbors and cells alike. Hence, the bullseye approach intuitively visualizes and quantifies the effect size, making it suitable for downstream analysis, such as comparing cross-expression between different regions.

We next conducted brain-wide analysis and found that 20% of possible ligand-receptor gene pairs and 4% of non-signaling gene pairs are cross-expressed, thus generating novel candidates that potentially encode functionally relevant interactions. In fact, these patterns are spatially enriched, where most gene pairs cross-express in a few slices and some cross-express in multiple slices (Extended Data Fig. 1a). We now highlight some notable examples of cross-expression for both signaling and non-signaling genes. The dopamine receptor D_1_ (*Drd1*) and proenkephalin (*Penk*) are strongly cross-expressed (Extended Data Fig. 1b), with discernible spatial enrichment in the striatal regions. *Drd1* is involved in the reward system [36,37] while *Penk* generates opioids that modulate fear response [38] and nociception [39,40], suggesting that these genes may be involved in avoidance behavior. Indeed, Penk is strongly co-expressed with the dopamine receptor D_2_ (*Drd2*) (Pearson’s *R* = 0.72 in scRNA-seq striatal data; *Drd2* is not in our gene panel), indicating that the D1 and D2 neurons are spatially intermingled, allowing them to play interrelated roles in motor control [41]. Additionally, we find that the somatostatin receptor *Sstr2* cross-expresses with vasoactive intestinal polypeptide receptor 1 (*Vipr1*/*VPAC1*) in the cortex (Extended Data Fig. 1c), suggesting a potential complementary interaction in modulating local neuronal circuits and influencing neuroendocrine signaling pathways [42]. Beyond the signaling genes, we note that the fibril-associated *Col19a1* (collagen type XIX alpha 1 chain), a gene involved in maintaining the extracellular matrix (ECM) integrity [43,44], cross-expresses with *C1ql3* (complement C1q like 3) (Extended Data Fig. 1d), whose secretion in the ECM facilitates synapse homeostasis and the formation of cell-cell adhesion complexes [45,46]. Finally, our analysis reveals that *Marcksl1* (myristoylated alanine-rich C-kinase substrate), which is involved in adherens junctions and cytoskeletal processes [47,48], cross-expresses with actin beta (*Actb*) (Extended Data Fig. 1e), hinting at their involvement in local tissue architecture [49]. Taken together, the cross-expression analysis not only reveals expected relationships between signaling molecules, but it also discovers genes implicated in the tissue microenvironment. Accordingly, cross-expression is an unbiased, data-driven framework for finding genes with orchestrated spatial expression profiles, with potential for novel discovery increasing with varying sizes and compositions of the gene panels.

We have thus far investigated cross-expression between cells and their neighbors. Yet, gene expression may be coordinated between more distant neighbors or between large spatial niches. The former is facilitated by changing the rank of the nearest neighbor tested. The latter is enabled by smoothing a gene’s expression in a cell by averaging it with its expression in nearby cells, as shown for cortical layer 4 marker *Rorb* (Fig. 2e) and layer 6 marker *Foxp2* (Extended Data Fig. 1f) in the auditory cortex [32].

Although cross-expression may appear at varying length scales, we focus our analyses at the cellular level to investigate its signature between individual cells.

### Cross-expression is driven by subtle and consistent cell subtype compositional differences

Having seen that cross-expression recovers coordinated spatial gene expression, we now explore its relationship with cell type heterogeneity. For this purpose, we use another BARseq dataset [35] that was recently used to create a mouse cortical cell type atlas using the same 104 excitatory marker genes as before. Here, we observe that genes cross-express between cells of the same and of different types. For example, *Gfra1* and *Foxp2* are cross-expressed within the same cell type L4/5 IT (intratelencephalic) and between different cell types Car3 or CT (corticothalamic) and L4/5 IT (Fig. 3a). In general, genes vary greatly in terms of the cell type labels of cross-expressing cell pairs (Fig. 3b). For instance, for some gene pairs 40% of the cell pairs have the same cell type label while in others as many as 90% of the cell pairs belong to different cell types (Extended Data Fig. 2a). Moreover, some genes involve many while others involve few cross-expressing cells. For example, in the analyzed data the median number of cross-expressing cell pairs is 2,378, and 27% of genes involve over 4,000 while only 5% involve 400 or fewer pairs (Extended Data Fig. 2b), indicating that the density of gene cross-expression is highly variable. Interestingly, cell type purity – the proportion of cell pairs with the same type – decreases as more cell pairs cross-express (Fig. 3c, Spearman’s *ρ* = – 0.46), highlighting a potential role for spatially intermingled cell types in patterns of cross-expression.

**Fig. 3:**
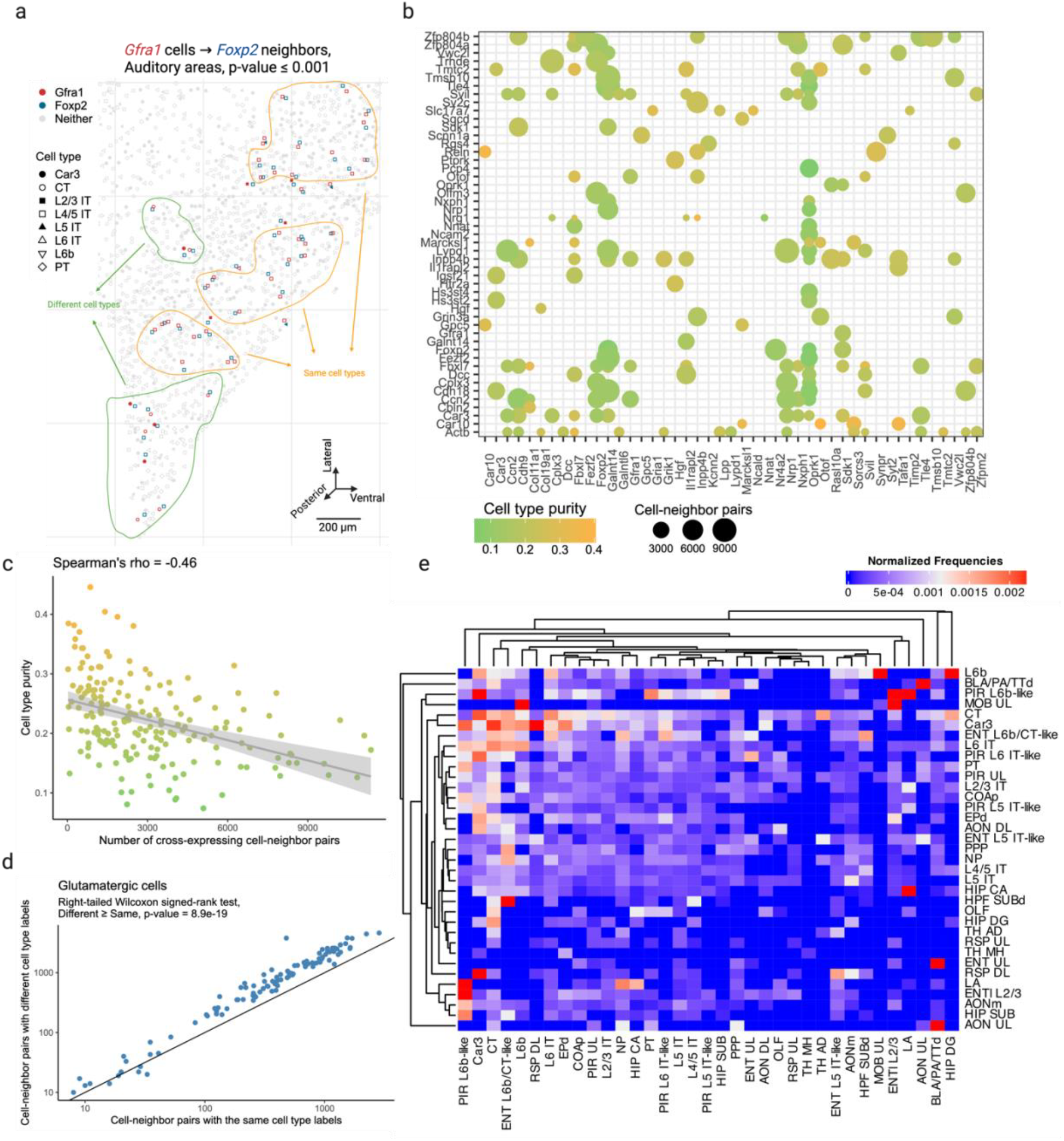
Cross-expression patterns are discovered independently of cell type labels but are driven by cell type heterogeneity. **a**, Cells of the same (yellow) and different (green) types cross-express genes *Gfra1* and *Foxp2* in the auditory cortex. Discovering cross-expression relations between this or any other gene pair does not require cell type labels. **b**, Genes are cross-expressed across numerous cells, with the dot size indicating the number of cell-neighbor pairs and the color showing the proportion of pairs with the same label (cell type purity). **c**, Cell type purity against the number of cross-expressing cell-neighbor pairs. Each point is a gene pair from (b), and shaded area is 95% confidence interval. **d**, Number of cell-neighbor pairs with the same or different cell subtype labels given that they were both labeled ‘glutamatergic’ at the higher level of the cell type hierarchy. Each point is a cross-expressing gene pair. **e**, Heatmap showing the normalized frequencies of cell type label combinations between cross-expressing cells. Created with BioRender.com.

To assess the influence of spatial cell type composition more broadly, we use our hierarchical cell type taxonomy [35], where types at a higher-level divide into subtypes at a lower level. Using cross-expressing glutamatergic cells, we find that 64% of the pairs consist of different cell subtypes (Fig. 3d, right-tailed Wilcoxon signed-rank test, different labels ≥ same labels, p-value ≤ 0.0001, Extended Data Fig. 2c), suggesting that subtle cell type differences drive cross-expression. However, for cross-expressing GABAergic cells, we find that only 44% of the pairs have different cell subtype labels (Extended Data Fig. 2d-e right-tailed Wilcoxon signed-rank test, different labels ≥ same labels, p-value = 1), reflecting the fact that our gene panel is optimized to detect cell subtype differences between excitatory, but not inhibitory, neurons. Crucially, we observe that cells of one type consistently cross-express with cells of another type (Fig. 3e, Extended Data Fig. 2f), indicating that cross-expression recapitulates patterns of cell type composition. Since cell type labels are assigned based on the expression of many genes, repeated spatial proximity of cell types results in the cross-expression of their marker genes.

### Cross-expression offers a common framework for analyzing multiple studies and the meta-analytic network reveals uniquely spatial biological processes

After analyzing ligand-receptor genes and cell type variability in terms of cross-expression, we asked whether its gene-centric approach can be used to simultaneously analyze multiple studies. To this end, we downloaded 13 adult mouse brain datasets (see methods), which were collected using 8 different platforms (4 commercial and 4 laboratory-based), spanning ~25 million cells (after dataset-specific quality control; median 724,790 cells) across 695 large samples/ slices (hemi-coronal, coronal, or sagittal) (Fig. 4a). Although the gene panels had varying sizes and compositions, they were sufficiently overlapping to allow integrative analyses (Fig. 4b; median pairwise overlap is 77 genes), reflecting that well-studied and informative genes are enriched across panels [50].

**Fig. 4:**
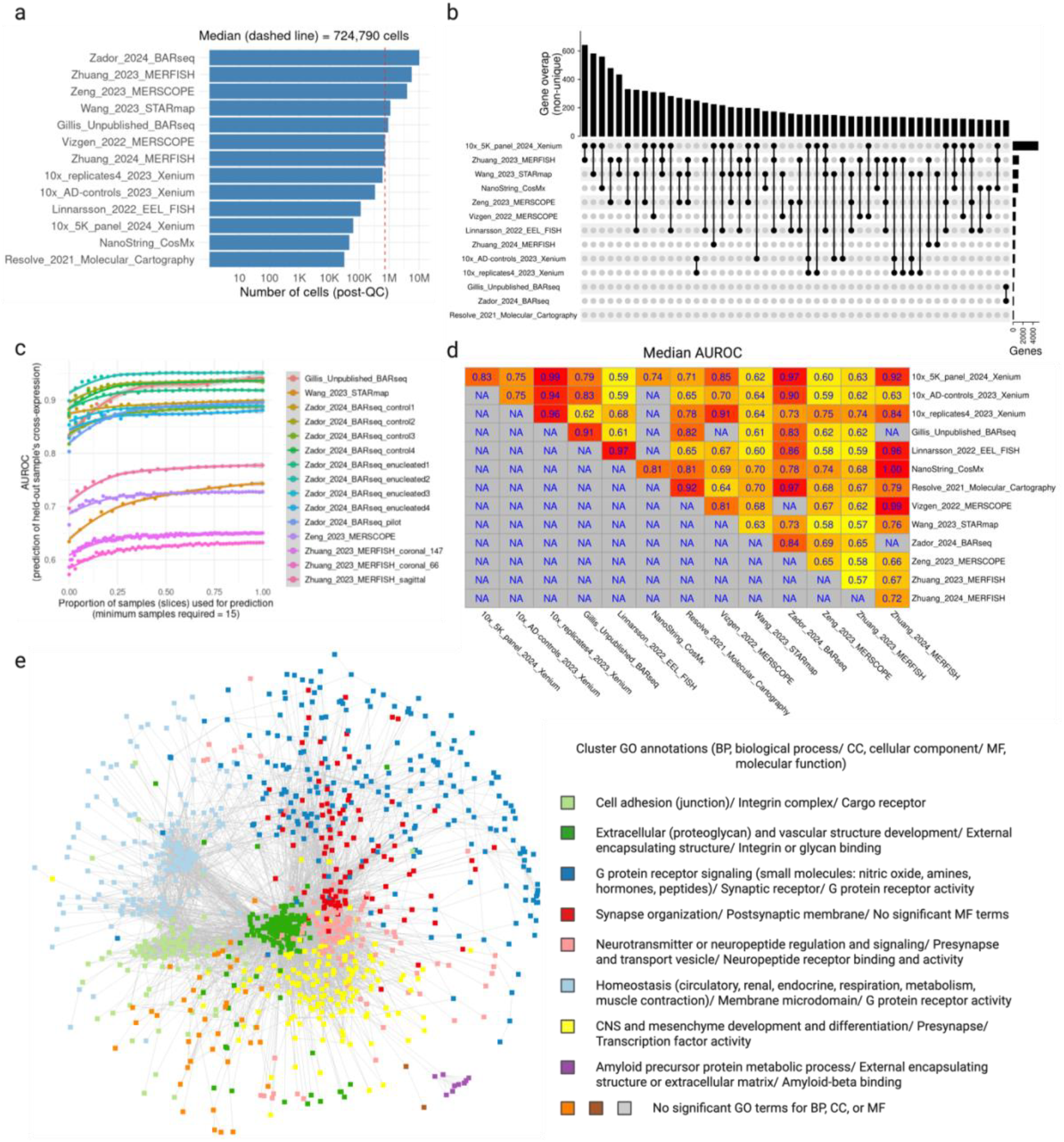
Cross-expression facilitates integrative meta-analysis of numerous samples across 13 studies. **a**, Number of cells in each dataset after applying dataset-specific quality control metrics. Red line is the median. **b**, Upset plot showing the overlaps between the gene panels of different datasets. **c**, Predictions of held-out slices’ p-values (significant or non-significant) using the average correlation of the remaining slices. Performance (AUROC) is reported as the function of the number of slices used to average the correlation (cross-validation). **d**, Median performance (AUROC) when predicting the p-values of a slice in one dataset using the correlation of slice(s) from another dataset using the genes shared between their panels. **e**, Cross-expression meta-analytic network where the nodes are genes, with edges present if the genes are cross-expressed in two or more datasets. The network is clustered into communities, with the colors and labels indicating gene ontology (GO) annotations, where slash (‘/’) separates biological process (BP), cellular components (CC), and molecular function (MF). Created with BioRender.com.

We first investigated cross-expression replicability within datasets. Here, we predicted the p-values of a held-out slice using the averaged correlations of the remaining slices. Our cross-validation-based classification revealed that the predictions (AUROCs) improved and then plateaued as more samples were included (Fig. 4c), suggesting that the sample ensemble progressively became more representative, e.g., due to the inclusion of anatomically adjacent slices. While the improvements were consistent across datasets, the AUROC scores varied between them, indicating the presence of potential batch effects, including differences in data quality between samples, enrichment of spatially variable or localized genes in the panels, and large gaps between the slices, etc. In general, the cross-expression patterns within datasets are broadly replicable, where including more samples improves performance, possibly due to better representation of the underlying anatomical regions.

Next, we examined whether cross-expression correlations in one dataset can predict the p-values in another dataset using the genes shared between their panels. We found high replicability for some datasets and modest performance for others (Fig. 4d), indicating that integrative meta-analysis is possible using at least a subset of these data. Although our results here are reported as median AUROCs, the performance can suffer if (say) in one dataset most slices are anterior while in the other most are posterior. Consequently, our subsequent analyses focus on replicability of cross-expression between (anatomically homologous) slices across datasets.

Finding that cross-expression is broadly replicable between datasets, we created a meta-analytic network, where the nodes represent genes and the edges indicate cross-expression (Fig. 4e). Our network includes gene pairs only when they are cross-expressed in two or more datasets, thus excluding those cross-expressed in just one dataset. The network shows high modularity, with communities enriched for gene ontology (GO) terms representing spatially-mediated biological processes. For example, the community ‘synapse organization’ (red) is connected with the ‘neurotransmitter regulation’ community (pink), recapitulating ligand-receptor interactions present in these data. Likewise, the ‘cell adhesion’ community (light green) is connected to the ‘extracellular and vascular’ community (dark green), implicating genes potentially involved in structural and supportive processes. The ‘G protein coupled receptor signaling’ community (dark blue) is diffusely connected to multiple other communities, reflecting these genes’ diverse physiological roles, such as in hormonal and metabolic processes. Similarly, the ‘CNS development’ community shows distributed connectivity, likely revealing the cross-expression of cell type markers with multiple other marker and non-marker genes, e.g., due to transcription factors (indirectly) specifying cell type identity [51]. Lastly, the network contains a small ‘amyloid precursor protein’ community (purple) composed of Alzheimer’s disease (AD) genes cross-expressed with *ApoE*, which helps form amyloid plaques and neurofibrillary tangles [52]. Together, the cross-expression framework allows us to use the shared genes to perform integrative analyses across multiple studies, with our meta-analytic network highlighting genes involved in distinctively spatial biological processes.

### Parkinson’s disease (PD) genes *Drd1* and *Gpr6* show highly reproducible and localized cross-expression patterns in the striatum

Our meta-analytic network revealed that cross-expressing genes are involved in spatially-mediated biological processes. Since genes were included in the network if they cross-expressed in two or more datasets, we sought to identify an example gene pair that is both highly reproducible and biologically informative. This process is inherently dependent on the researchers’ interests, so we focused on *Drd1* and *Gpr6*, which are implicated in Parkinson’s disease (PD) due to their hypo- and hyperactivity, respectively [53–60]. Our datasets are registered to the Allen Institute’s CCFv3 mouse brain region atlas [61], which allowed us to arrange samples from multiple brains and different datasets in the same anatomical space. Looking at coronal and sagittal sections, we found these genes’ cross-expression highly concentrated within a restricted range and effectively absent outside it, suggesting that it is localized within specific anatomical regions (Fig. 5a). Next, we overlaid the Allen CCFv3 atlas on the combined cross-expression signal, finding that it is located within the striatum (Fig. 5b, protruded region), the central locus in PD pathophysiology [62]. Consistent with this observation, we found that their cross-expression within slices is located in the striatal regions but not elsewhere, with its patterns following gross neuroanatomy in the anterior-to-posterior and lateral-to-medial directions (Fig. 5b, panels 1-9 and P-S). Accordingly, the genes *Drd1* and *Gpr6* show coordinated expression between cells in the striatum, a property observed in multiple slices across different datasets.

**Fig. 5:**
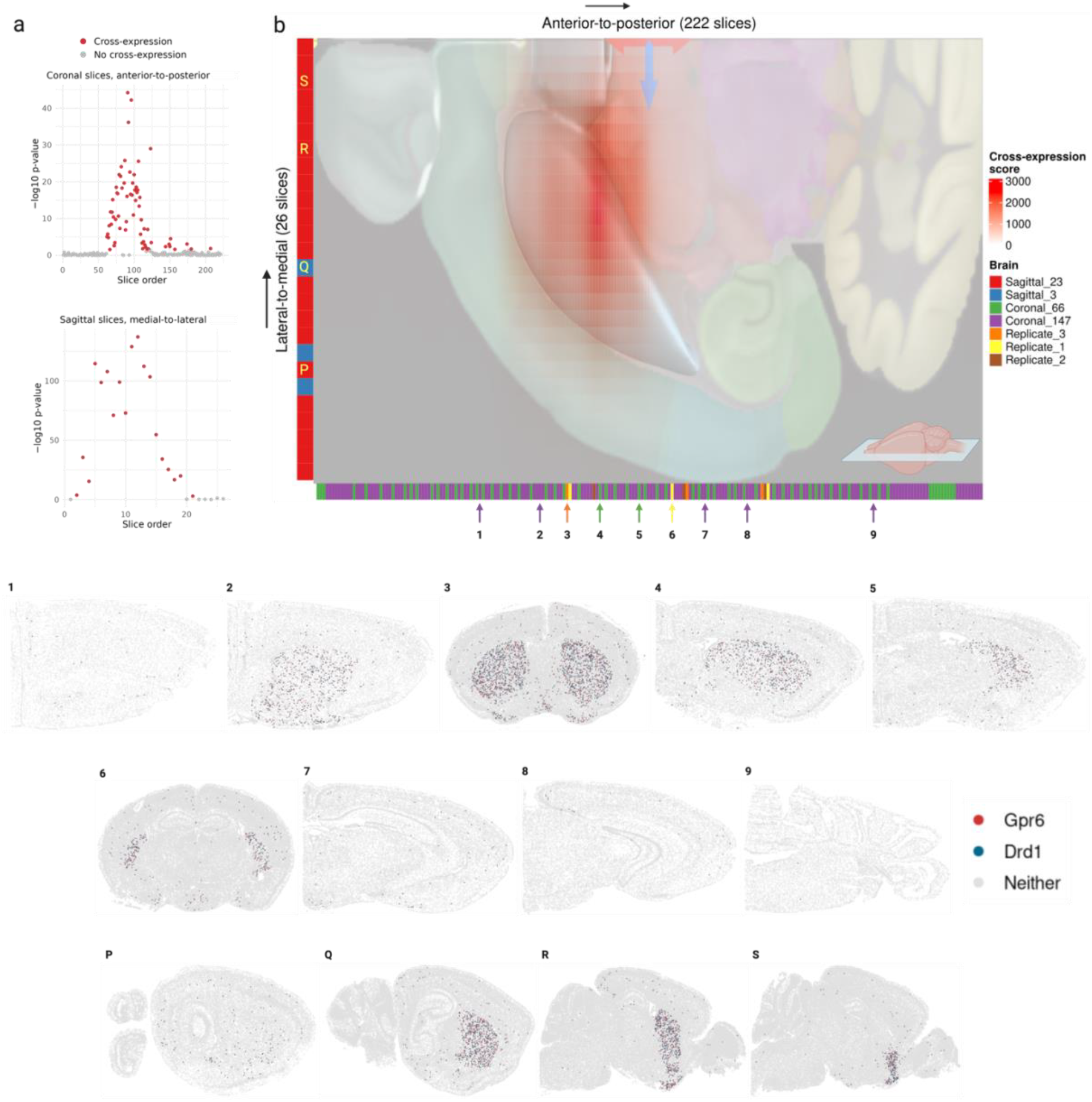
Cross-expression of genes *Drd1* and *Gpr6* in the striatum across multiple brains and different studies. **a**, Cross-expression signature (–log10 p-value) of *Drd1* and *Gpr6* genes in the coronal sections ordered in the anterior-to-posterior direction using the Allen Institute CCFv3 mouse brain atlas coordinates (top). Bottom, same as top but with sagittal slices ordered in the medial-to-lateral direction. **b**, Cross-expression score (outer product of coronal and sagittal slices’ –log10 p-values) overlaid with the Allen Institute CCFv3 brain region annotations, with the striatum as the protruded (blue) area. Different colors represent the source brain of each slice. Numbers (1-9) and letters (P-S) indicate the position of example coronal and sagittal slices, respectively, within the overall gross neuroanatomy. Created with BioRender.com.

The cross-expression of these genes can help facilitate subsequent studies. Briefly, *Drd1* is the dopamine receptor D1, whose stimulation by dopamine initiates movement, whereas *Gpr6* is the G protein coupled receptor 6, whose constitutive activity prevents movement initiation. In PD, the dopaminergic neurons in the substantia nigra insufficiently activate *Drd1*, and *Gpr6* exhibits higher baseline activity, leading to severe difficulty in starting movement, the cardinal symptom of PD [53–60]. While extant therapies increase dopamine (L-DOPA as Levodopa) to stimulate *Drd1* [63,64], where the drug’s effectiveness decreases over time and causes sides effects, recent clinical trials have explored the inverse agonist CVN424 to inhibit *Gpr6* [55–60]. Although at present these approaches have not been pursued in tandem, their localized cross-expression suggests that the drugs’ staggered or co-delivery might offer complementary, potentially amplified, effects. In general, once reproducible cross-expression signatures are discovered, ideally across different samples and studies, one can further investigate gene pairs of interest, making cross-expression a basic tool for biological research using spatial transcriptomic data.

### Cross-expression network reveals *Gpr20* as a central gene and discovers possible interaction partners between astrocytes and the brain microvasculature

Having assessed multiple datasets, we now analyze a single study to show how cross-expression patterns can be used alongside additional information, such as cell type labels. We first supplemented our network formalism by including (second-order) edges between two genes if they independently cross-expressed with a third same gene (Fig. 6a). Using the Vizgen MERFISH data, we created a cross-expression network (Fig. 6b), which contains 200 genes with 382 first-order, 217 second-order, and 107 dual-order edges. We observe that *Gpr20*, a G protein-coupled receptor, is a central gene with a high node degree of 40 while the other genes form a median of 4.8 edges (Extended Data Fig. 4a). We performed gene ontology (GO) enrichment for genes cross-expressed with *Gpr20*, finding functional groups like ‘regulation of macromolecule biosynthetic process’, ‘regulation of gene expression’, and ‘regulation of metabolic process’ (Extended Data Fig. 4b, all p-values ≤ 0.05). While some of these genes are co-expressed with astrocytic and microglial cell type markers (Extended Data Fig. 4c), their global co-expression with the endothelial marker is higher, where the co-expression profiles were computed using neighbors cross-expressed with *Gpr20* rather than the entire dataset (Extended Data Fig. 4d, Mann-Whitney U test, endothelial vs others, all p-values ≤ 0.01; remaining pairwise comparisons, all p-values > 0.05).

**Fig. 6:**
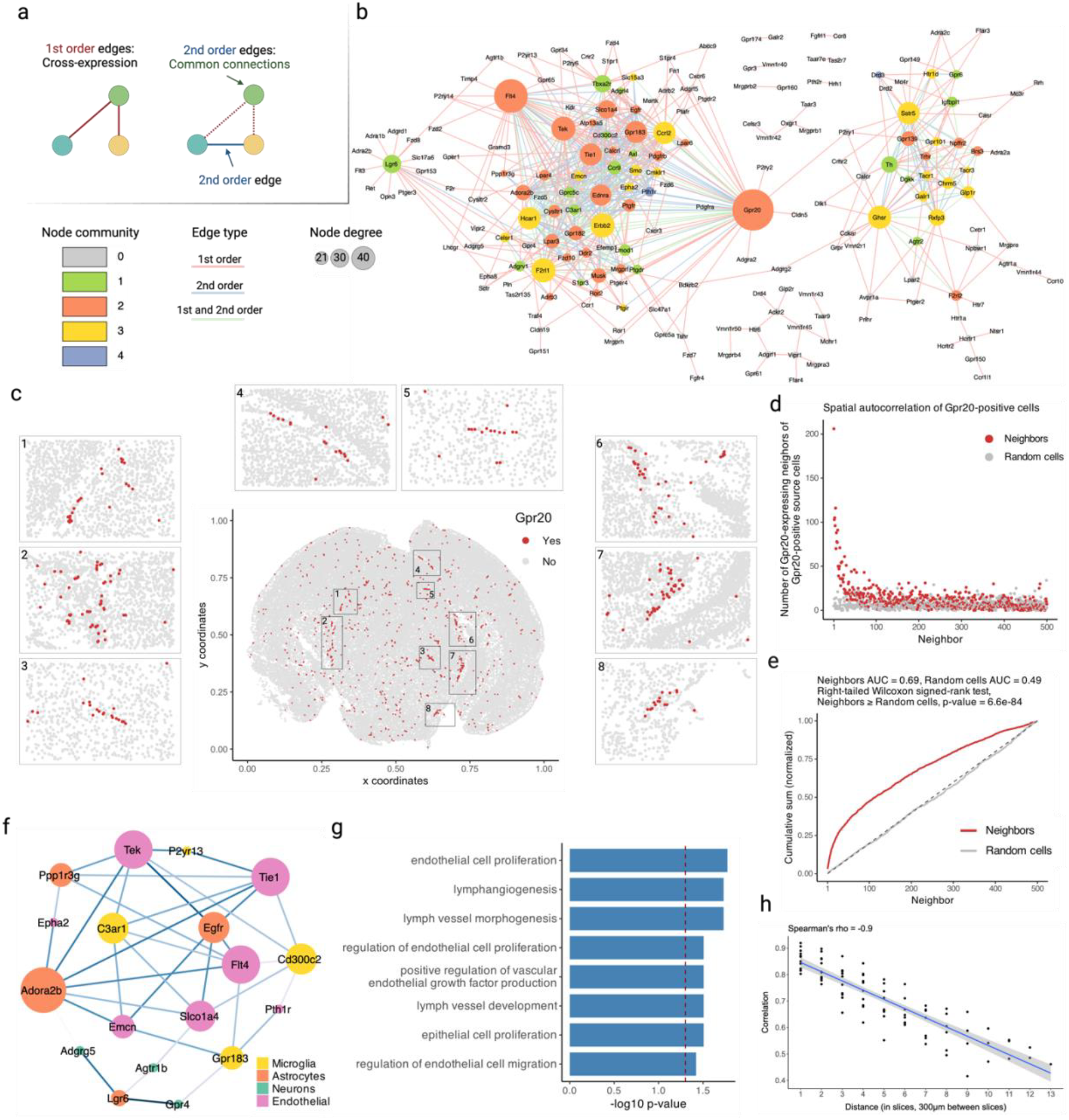
Networks of cross-expression. **a**, Cross-expression (edges) between genes (nodes) forms a network (left), where second-order edges (right) between genes share a node (first-order). **b**, Example cross-expression network, with first-order node degree represented by size and edge color showing first-, second-, or dual-order (first-order and second-order) connections. Threshold for the second-order edges is 4. Node color shows community membership assigned by Louvain clustering the second-order network. **c**, Cells are colored based on *Gpr20* expression. Numbered rectangles in the central figure correspond to zoomed-in versions. **d**, Number of neighbors with *Gpr20* given that the source cells also express this gene. **e**, Cumulative sums (after L1 normalization) from (d) for true and randomly selected neighbors. Identity line is dashed. **f**, Subnetwork created from (b) by pruning edges with significant co-expression and then removing nodes with degree 1. Nodes are colored by cell types based on their co-expression with marker genes. **g**, GO functional groups for genes in the subnetwork in (f). **h**, Similarity in the network structures of nearby and distant brain slices. Shaded area is 95% confidence interval. Created with BioRender.com.

Noting that the neighbors of *Gpr20*-positive cells are involved in the microvasculature, we next viewed the spatial distribution of cells expressing *Gpr20*, finding that they form contiguous linear streaks resembling blood vessels (Fig. 6c; anterior slice from mouse brain 1 shown). To test this observation, we looked at whether the neighbors of *Gpr20*-positive cells also express this gene and compared it to randomly selected cells, which constitute the expectation that *Gpr20* is uniformly expressed across space. Consistent with the visualization, we find that cells with *Gpr20* are surrounded by neighbors that also express this gene, a pattern that disappears for neighbor order of 50 or more cells (Fig. 6d-e, area under curve (AUC), neighbors vs random cells, 0.69 vs 0.49; right-tailed Wilcoxon signed-rank test, neighbors vs random cells, p-value ≤ 0.0001). Having seen that cells with *Gpr20* possibly reflect blood vessels, we asked whether these cells are themselves vascular or whether they line the vasculature, especially since the cells that cross-express with *Gpr20* are endothelial. We find *Gpr20* is poorly co-expressed with *Igfr1* (Pearson’s *R* = 0.0024), the vascular/endothelial marker [65–67] in our gene panel, suggesting that it lines but does not mark the blood vessels. Moreover, it is lowly co-expressed with other cell type markers (average Pearson’s *R*, astrocytes = –0.0027, microglia = 0.0018, oligodendrocytes = – 0.022, neurons = –0.0025), eschewing cell type characterization. Taken together, *Gpr20*, a salient topological feature of our cross-expression network, seems to be expressed in diverse cell types that line the blood vessels, reflecting its possible role in the microvasculature.

Cross-expression driven by cell types might be particularly common when two genes which cross-express with a third gene are co-expressed together, reflecting some common transcriptional program jointly cross-expressing with neighboring cells. To investigate this, we reduced co-expression further by specifying that cross-expressing genes must show lack of significant co-expression, a procedure that yielded a subnetwork, which we further curated by removing genes with fewer than two edges. Indeed, we find that two genes that independently cross-express with another gene tend to be co-expressed (Fig. 5f, Extended Data Fig. 5a) and, as expected, belong to the same cell types, as revealed by their co-expression with cell type marker genes (Extended Data Fig. 5b). Confirming these results, the subnetwork genes are enriched in GO groups like ‘endothelial cell proliferation’, ‘positive regulation of vascular endothelial growth factor production’, and ‘regulation of endothelial cell migration’ (Fig. 6g, all p-values ≤ 0.05). These results indicate that while cross-expressing genes are present in specific cell types, the relations between them are functionally suggestive as opposed to simply reflecting cell type compositional differences, especially since the cell type markers are not cross-expressed. For example, the astrocytic *EGFR* (epidermal growth factor receptor) cross-expresses with the vascular *Flt4*/*VEGFR-3* (FMS-like tyrosine kinase 4), *Tek*/*Tie2* (TEK tyrosine kinase/ angiopoietin-1), and *Tie1* (tyrosine kinase with immunoglobulin-like and EGF-like domain 1). These three vascular receptors promote angiogenesis via the *VEGF* (vascular epidermal growth factor) ligand [68,69], prevent endothelial cell apoptosis [70,71], and negatively regulate angiogenesis [72], respectively, thus reflecting their potential role in the brain microvasculature in coordination with the astrocytes, whose endfeet ensheathe the blood microvessels to constitute the blood-brain barrier (BBB).

Within the same subnetwork, the astrocytic gene *Ppp1r3g* (protein phosphatase 1 regulatory subunit 3G), which helps convert glucose to glycogen [73], cross-expresses with *Epha2* (ephrin type-A receptor 2), whose activity makes the BBB more permeable [74], likely enabling glucose’s transport and eventual conversion into glycogen, thereby making this cross-expression relation relevant for energy metabolism. Indeed, this observation can be used to generate hypotheses about the (directional) relationship between energy needs within a local microenvironment and remodeling of the microvasculature, making cross-expression a powerful approach with which to form testable hypotheses.

Next, we asked whether cross-expression networks change across the brain. Because gene expression is regional, slices from various areas should show cross-expression between distinct genes. We assessed this by forming networks for each slice in our sagittal BARseq data. As expected, we find that adjacent slices have similar networks than distant slices (Fig. 6h, Spearman’s *ρ* = –0.9), a trend also seen in our BARseq coronal data (Extended Data Fig. 6a, Spearman’s *ρ* = –0.87) but not when the two datasets are mixed and the “distance” reflects the difference in the order of slices (Extended Data Fig. 6b, Spearman’s *ρ* = 0.094). Hence, cross-expression is sensitive to broad spatial variation in gene expression.

### Cross-expression discovers anatomical marker gene combinations that delineate the thalamus and refine cortical layer VI boundaries

A key goal in biological research is finding marker genes, such as those that identify cell types, e.g., *Olig1* for oligodendrocytes [32], anatomical regions, e.g., *Rorb* for cortical layer IV [35], and functional modules, e.g., *Trpc4* for lateral septum in social behaviors [75]. In addition to using individual markers, one can use co-expression to discover marker gene combinations, an approach that uses single-cell or single-nucleus RNA-seq databases [76]. Although co-expression provides more combinations than individual markers, it relies on measurements from the same cells, thereby underutilizing the spatial relations between cells in spatial transcriptomics datasets. Leveraging the spatial dimension to discover marker gene combinations, we asked whether cross-expressing genes can delineate anatomical regions, including putative functional modules. As an example, we found that cross-expression between *Lgr6* and *Adra2b* delineates the thalamus even though these genes are expressed throughout the brain (Fig. 7a). Specifically, while 48% of *Lgr6*- and 57% of *Adra2b*-expressing cells are thalamic, 91% of their cross-expressing cell-neighbor pairs are in the thalamus (Extended Data Fig. 3a), underscoring the spatial enrichment of their cross-expression signature (Extended Data Fig. 3b). We find that *Lgr6* also cross-expresses with *Ret* in the thalamus despite brain-wide expression of both genes (Fig. 7b, Extended Data Fig. 3c). Next, we examined whether *Adra2b* and *Ret*, both of which cross-express with *Lgr6*, show enriched co-expression in the thalamus. We find that they are indeed co-expressed within the thalamus but not in the rest of the brain (Fig. 7c), e.g., 89% of their co-expressing cells are in the thalamus, thus serving as robust combinatorial markers.

**Fig. 7:**
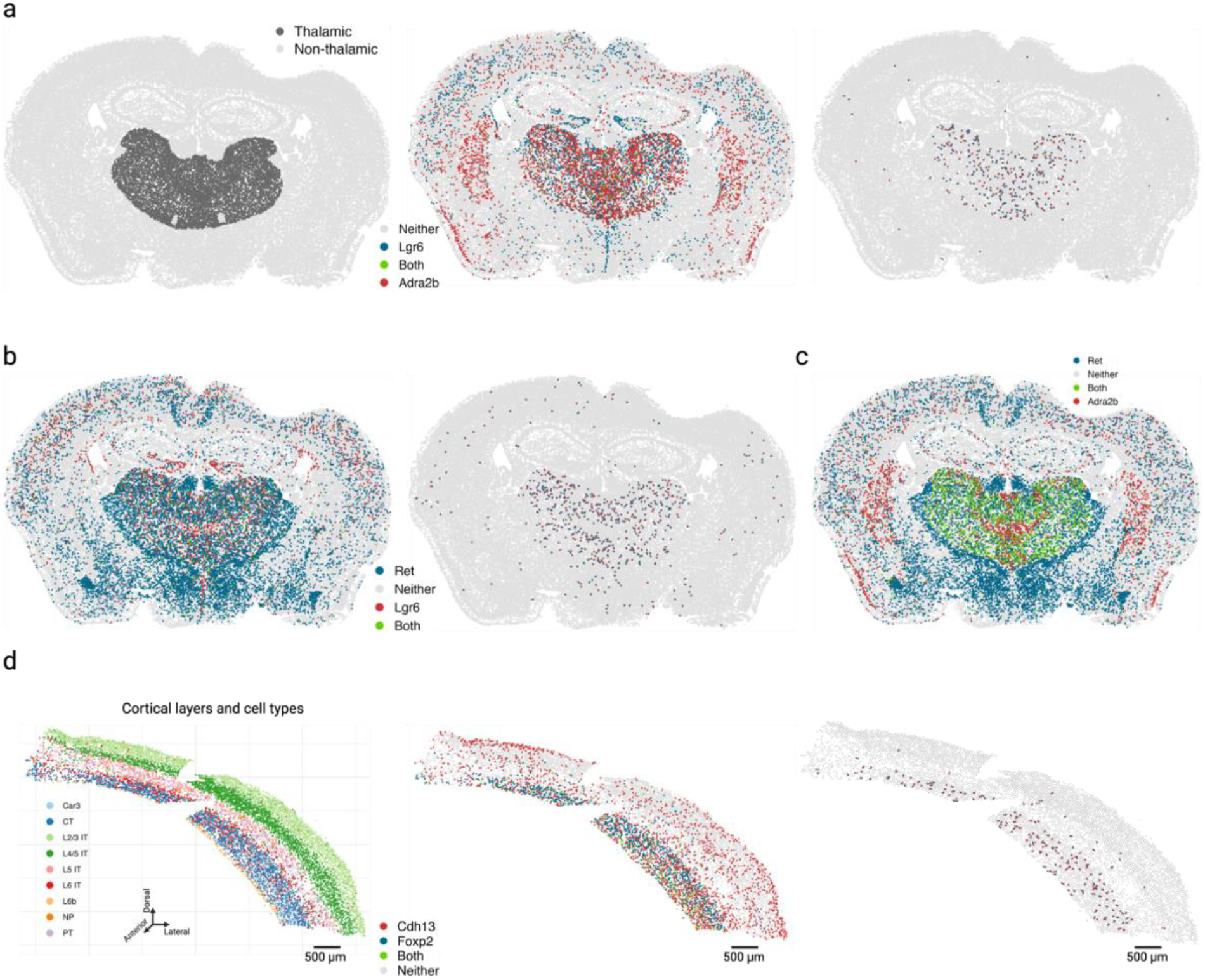
Cross-expression can discover combinatorial anatomical markers. **a**, Comparing the thalamus (left) to the rest of the brain, genes *Lgr6* and *Adra2b* are widely expressed across multiple brain regions (middle) but are preferentially cross-expressed in the thalamus (right). **b**, Same as in (a) but for genes *Lgr6* and *Ret*. **c**, Genes cross-expressing with *Lgr6* in (a) and (b) co-express in the thalamus. **d**, Cross-expression of *Cdh13* with cortical layer 6 marker *Foxp2* (middle) recapitulates layer 6 boundaries (right, cf. left). Created with BioRender.com.

To evaluate whether the combinatorial marker-based approach is reliable, we asked whether single gene markers, when assessed for cross-expression, rediscover the anatomical locations. Using the BARseq cortical cell type atlas data [35], we assessed cross-expression between cortical layer 6 marker *Foxp2* and ubiquitously expressed gene *Cdh13*. We discover that cross-expression between these genes delineates layer 6 boundary (Fig. 7d), further supporting the view that combinatorial anatomical markers can be discovered using cross-expression. Indeed, the layer 6 boundary recovered by cross-expression captures additional L6 IT neurons whereas *Foxp2*-based boundary overlooks these cells, indicating that combinatorial markers can refine extant anatomical regions. More generally, this process leverages the spatial enrichment of cross-expression, where we assess whether cross-expressing pairs are closer to other cross-expressing pairs than to randomly selected cells. Our framework therefore discovers gene pairs that annotate anatomical regions and refine extant boundaries, thus extending the single gene and co-expression-based approaches.

### Cross-expression signal is replicable across datasets, and global co-expression between spatial and single cell datasets indicates reliable cell segmentation

Two sources of non-biological variation in spatial transcriptomics [2–8] are batch effects, which result from technical differences between experimental runs, and cell segmentation, which draws boundaries around and assigns transcripts to cells, a process that can alter gene expression profiles and affect downstream analysis, including cross-expression.

We assess batch effects by comparing cross-expression between corresponding slices across biological replicates. The MERFISH data contains three replicates with three slices each, where the slices are sampled from roughly the same position. We find that the cross-expression signature is highly similar across replicates. For example, the average correlation for the anterior slices between the three replicates is 0.83 (Fig. 8a), with similar findings for the middle and posterior slices (Extended Data Fig. 7a-b, Spearman’s *ρ* = 0.81 and 0.8, respectively).

**Fig. 8:**
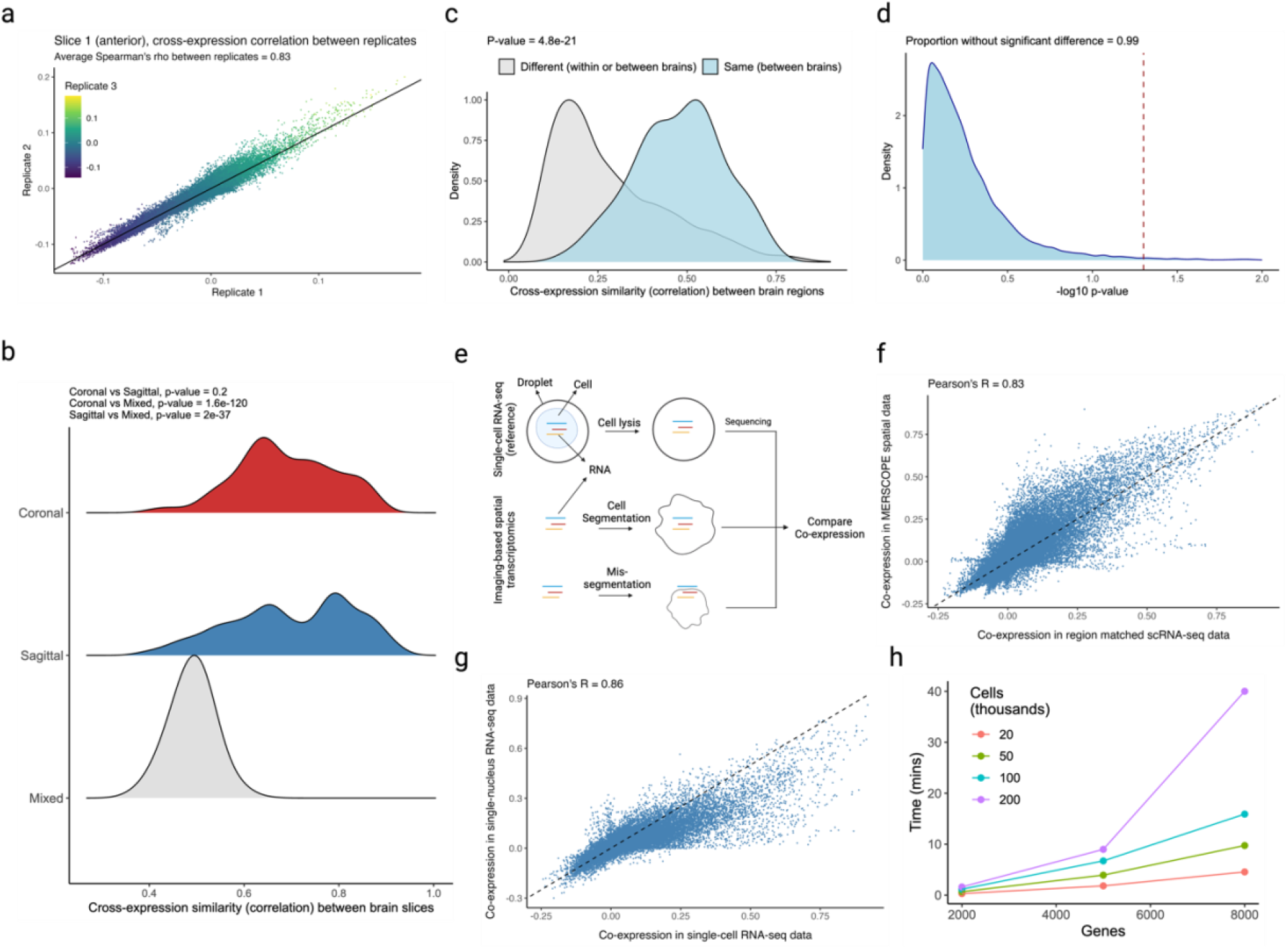
Assessing batch effects, cell segmentation, and software runtime. **a**, Correlation between cross-expression signatures across three biological replicates. **b**, Correlation between cross-expression signatures within (sagittal or coronal) and between (mixed) brains. Positive signal between brains likely reflects the fact that the sagittal and coronal brains both contain regions in the dorsal to ventral direction. **c**, Correlation between cross-expression signatures between the same anatomical regions across brains or between different anatomical regions across or within brains. **d**, Density of cross-expressing cells in the dorsal to ventral directions is compared across the sagittal and coronal brains. Significant p-values (without FDR correction) indicate that a cross-expressing gene pair has different densities across the two brains. Red dotted line is the significance threshold at alpha = 0.05. **e**, Single cell RNA-sequencing (scRNA-seq) profiles cells’ gene expression without cell segmentation. Co-expression between scRNA-seq and spatial transcriptomic data helps diagnose segmentation artifacts. **f**, Gene co-expression in spatial transcriptomic and in scRNA-seq data. Each point is a gene pair. **g**, Gene co-expression in single-nucleus RNA-sequencing (snRNA-seq) and in scRNA-seq data. Same gene panel is used in (f) and (g). **h**, Software runtime for varying numbers of genes and cells using a personal laptop with 16 GB RAM. Created with BioRender.com.

We next assessed the degree to which cross-expression within the BARseq sagittal or coronal experiments [35] is similar to that between experiments. To this end, we compared cross-expression patterns between brain slices. As expected, the cross-expression profiles are more similar within brains than between brains (Fig. 8b, Mann-Whitney U tests, FDR-corrected, coronal vs sagittal, p-value = 0.2, coronal vs mixed, p-value ≤ 0.001, and sagittal vs mixed, p-value ≤ 0.001), suggesting that the sectioning procedure samples different brain regions and therefore reveals distinct underlying gene expression profiles. Supporting this result, we find that the same anatomical regions (per Allen CCFv3 brain atlas [61]) across brains have more similar cross-expression profiles than do different regions within or between brains (Fig. 8c, Mann-Whitney U test, different regions vs same regions, p-value ≤ 0.001). Noting that the sagittal and coronal brains contain the same regions in the dorsal to ventral directions, we asked whether the cross-expression is similar in this shared dimension. Here, we computed the density of cross-expressing cells in the dorsal to ventral direction and compared these distributions across the two brains, finding that 99% (without FDR correction) of the genes did not have significantly different density profiles (Fig. 8d), suggesting that the cross-expression patterns are highly similar across batches at the whole-brain level.

Having found that the cross-expression profiles are generally robust, we assessed cell segmentation at a global level by comparing gene co-expression between the single cell RNA-sequencing (scRNA-seq) [32] and spatial transcriptomic data. We reasoned that scRNA-seq does not require segmentation and therefore captures genes co-expressed within the cell’s boundaries (Fig. 8e). Because cell segmentation alters transcript assignment, it could change co-expression in spatial transcriptomic data. Reassuringly, we find a strong association between gene co-expression in the scRNA-seq and spatial transcriptomic data (Fig. 8f, Pearson’s *R* = 0.83). We further examine whether this correlation is sufficiently strong by comparing co-expression between scRNA-seq and single-nucleus RNA-sequencing (snRNA-seq) [77] (Fig. 8g, Pearson’s *R* = 0.86), finding agreement between the two comparisons (*R* = 0.83 vs. *R* = 0.86). These results imply similar levels of technical variability between platforms while suggesting that gene co-expression is congruent between scRNA-seq and spatial transcriptomic data.

The data in our work was processed using CellPose [78], a deep learning-based cell segmentation algorithm. A recent benchmarking study showed that it outperforms other methods on a variety of metrics [79]. In fact, it uses the nuclear stain DAPI as a cell landmark and forms boundaries using cytoplasmic signal, such as the transcript distributions, making it the state-of-the-art segmentation algorithm on a variety of assessments. Further, the cell segmentation algorithms are continuously being improved [80], allowing users to re-segment and reanalyze their data. Most importantly, the analysis conducted using the cross-expression framework may suffer if segmentation is performed poorly, but the validity of the concept and the soundness of its statistics do not rely on this potential artefact and, with rapid improvements in data quality, the inferences drawn from it will become increasingly more reliable.

Moreover, we assessed the relationship between cross-expression and noise in gene expression measurement. Since the algorithm requires binarizing the expression matrix, an appropriate threshold needs to be applied prior to analysis. To count a gene as expressed in a cell, we applied thresholds of 1 to 10 molecules, finding that the cross-expression patterns are generally concordant across these noise levels (Extended Data Fig. 8a-b, median Pearson’s *R* = 0.88). Importantly, our framework is agnostic to and compatible with multiple models of gene expression noise [81], and once an appropriate threshold has been applied, the resultant expression matrix can be used for cross-expression analysis.

Finally, we explored the patterns of cell-neighbor relations and found that over 60% of cells are the nearest neighbors of exactly one cell but the remaining cells are the nearest neighbors of two or more cells (Extended Data Fig. 9a). Patterns such as these may be biologically important if the ‘neighbor’ cell plays a central role in the local microenvironment, so deviations from one-to-one mappings should be captured by statistical analyses. To investigate that our results are consistent across these patterns, we compared cross-expression in one-to-one against many-to-one mappings and with the full dataset, finding an average Pearson’s correlation of 0.96 (Extended Data Fig. 9b). Importantly, our procedure is consistent with the assumption of independent sampling because while a cell may be the nearest neighbor of multiple cells, each cell-neighbor pair is statistically independent.

We facilitate these analyses by providing a highly efficient R package. A laptop with 16 GB RAM can test for cross-expression in large datasets containing hundreds of thousands of cells and thousands of genes within minutes (Fig. 8h). At present, most (commercial) imaging-based platforms cannot profile gene panels of this magnitude [2–8], though such capabilities are anticipated. Our software’s performance makes it well-suited for analyzing current and future spatial transcriptomic datasets.

## Discussion

Cross-expression allows us to study gene-gene networks that reflect how nearby cells influence each other by coordinating their gene expression. Using this framework, we recapitulated known ligand-receptor interactions at the single cell level, revealing biologically meaningful tissue phenotypes. We further showed that cross-expression is discovered without cell type labels but often reflects cell subtype compositional differences in the form of marker gene cross-expression. We also revealed that its gene-centric perspective enables integrative meta-analysis, where many studies can be combined to find robust biological signals, such as the cross-expression of *Drd1* and *Gpr6* in the striatum. Moreover, it helps us perform deeper analyses of individual studies, revealing the relationships between astrocytes and the brain microvasculature, and discovers paired markers for anatomical region annotation. Together, cross-expression is a powerful way of analyzing spatial transcriptomic data and allows us to study gene-gene relations between adjacent cells, thereby fully harnessing the high-throughput of these technologies.

The cross-expression framework complements current approaches analyzing spatial transcriptomic data, such as those exploring niche-specific co-expression patterns [13–19]. Specifically, niche-specific cross-expression networks may be compared with co-expression networks to examine if inter-cellular relations are associated with intra-cellular gene programs and vice versa. This may be approached at different, potentially hierarchical spatial scales to reveal spatial gene expression programs within the tissue. Moreover, the cross-expression patterns can be quantified in multiple ways, such as using mutual information or graphlets, allowing investigations into the best approaches that capture the signal of interest. For example, just as co-expression relations can be measured using the Pearson’s correlation coefficient, cross-expression patterns may be investigated from numerous perspectives to discover the most robust formalism. In this sense, the cross-expression framework introduced here is primarily a way of conceptualizing gene-gene relations within spatial transcriptomics data, thereby serving as a powerful framework for research in tissue biology. For instance, it can be used to study cancer [82], where tumor progresses via signaling with the stromal tissue, as well as neurodegenerative diseases like Alzheimer’s [83] or senescence [84], where the progression of pathology is spatially structured, making it a broadly useful approach for numerous problems.

Cross-expression is not restricted to imaging-based spatial transcriptomics. Instead, it can be applied to any biological assay that provides cell-by-features and cell-by-coordinate matrices. For example, it can be extended to spatial proteomics [85], with potential to discover ligand-receptor interactions. Likewise, it may be applied to spatial translatomics [86] to focus on translating mRNAs that are more likely to form functional proteins, making conclusions about cell-cell relations more robust. In fact, with the increasing resolution of spatially barcoded RNA capture based methods [87,88], the framework may be extended transcriptome-wide to understand relations between spots at near single-cell resolution.

A key challenge in imaging-based spatial transcriptomics [2–8], including the datasets used in this work, is the size and constitution of the gene panels, which sets an upper limit on biological discovery. Although our framework will become more powerful as the quality of spatial transcriptomic data, especially the gene panel, increases, care must be taken to not interpret the results in mechanistic terms. Instead, the coordinated gene expression between neighboring cells should be viewed as a target for further investigation. In this sense, the cross-expression framework substantially narrows the space of gene-gene relations by identifying pairs that are potentially biologically meaningful, making the problem experimentally tractable. Overall, cross-expression offers a unique and powerful perspective on using spatial transcriptomic data for driving biological discovery.

## Conclusions

Cross-expression is a useful conceptual and analytical framework, which compares all genes and identifies pairs that coordinate their expression between neighboring cells. The accompanying R software efficiently facilitates these and other analyses as well as provides new set of visualizations to deeply explore spatial transcriptomics data by leveraging coordinated gene expression at the single-cell resolution.

The package is available at https://github.com/gillislab/CrossExpression.

## Extended Data Figures

**Extended Data Figure 1:**
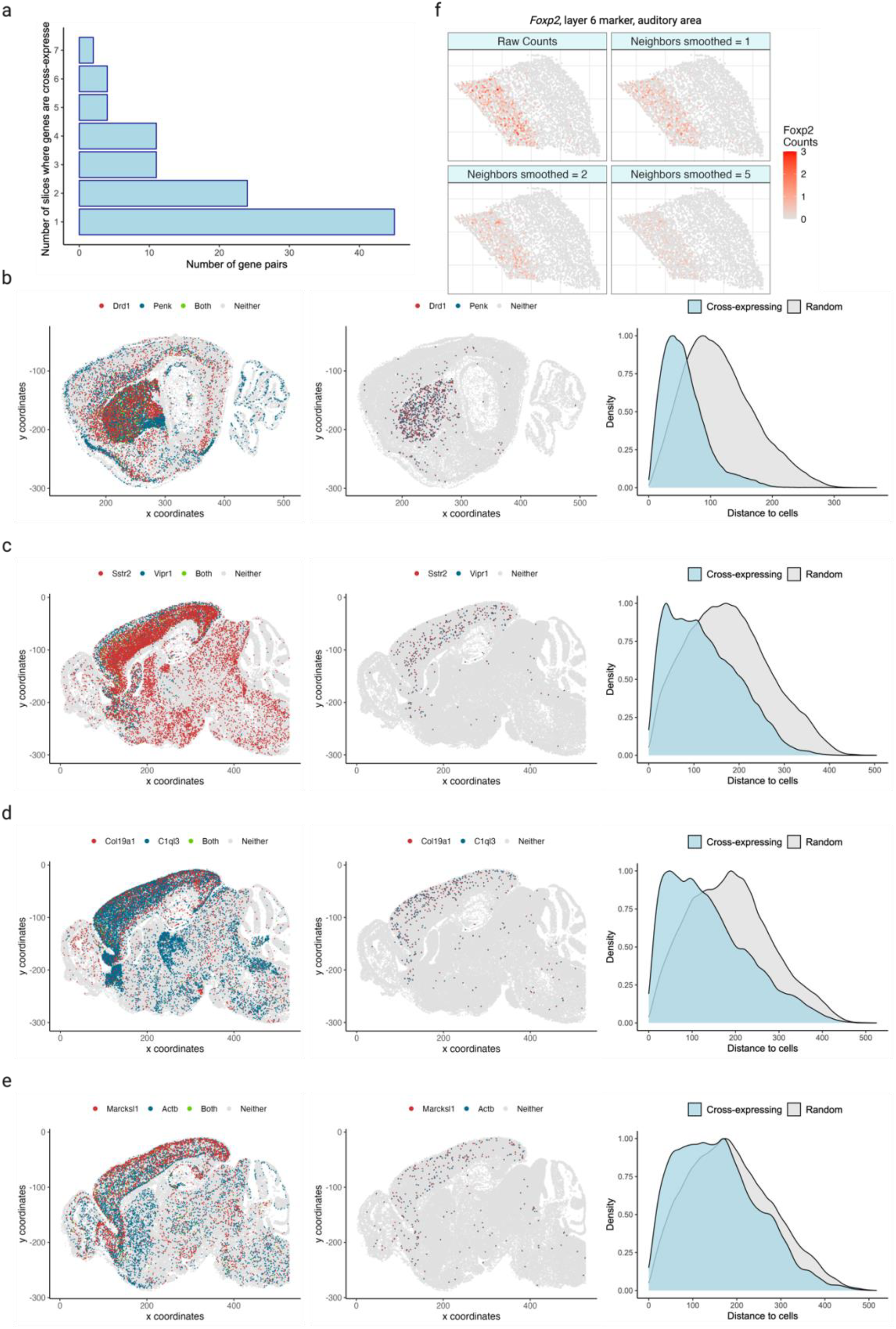
Cross-expression across tissue slices and regions for ligand-receptor and non-signaling genes. **a**, Distribution of the number of gene pairs cross-expressed in different slices. Dataset has 16 slices sampled sagittally from the left hemisphere of a mouse brain. **b-e**, Cells are colored by gene expression (left) and cross-expressing cells are highlighted (center). Right, distances between cross-expressing cells are compared with those between cross-expressing and randomly selected cells. Smaller distances mean that cross-expressing cells are nearer each other (spatial enrichment) than expected by chance (p-values ≤ 0.01, left-tailed Mann-Whitney U test). Genes include ligands and receptors (b, c) and non-ligands and non-receptors (d, e). **f**, Smoothed gene expression for different numbers of neighbors for the auditory cortical layer 6 marker gene *Foxp2*. Created with BioRender.com.

**Extended Data Figure 2:**
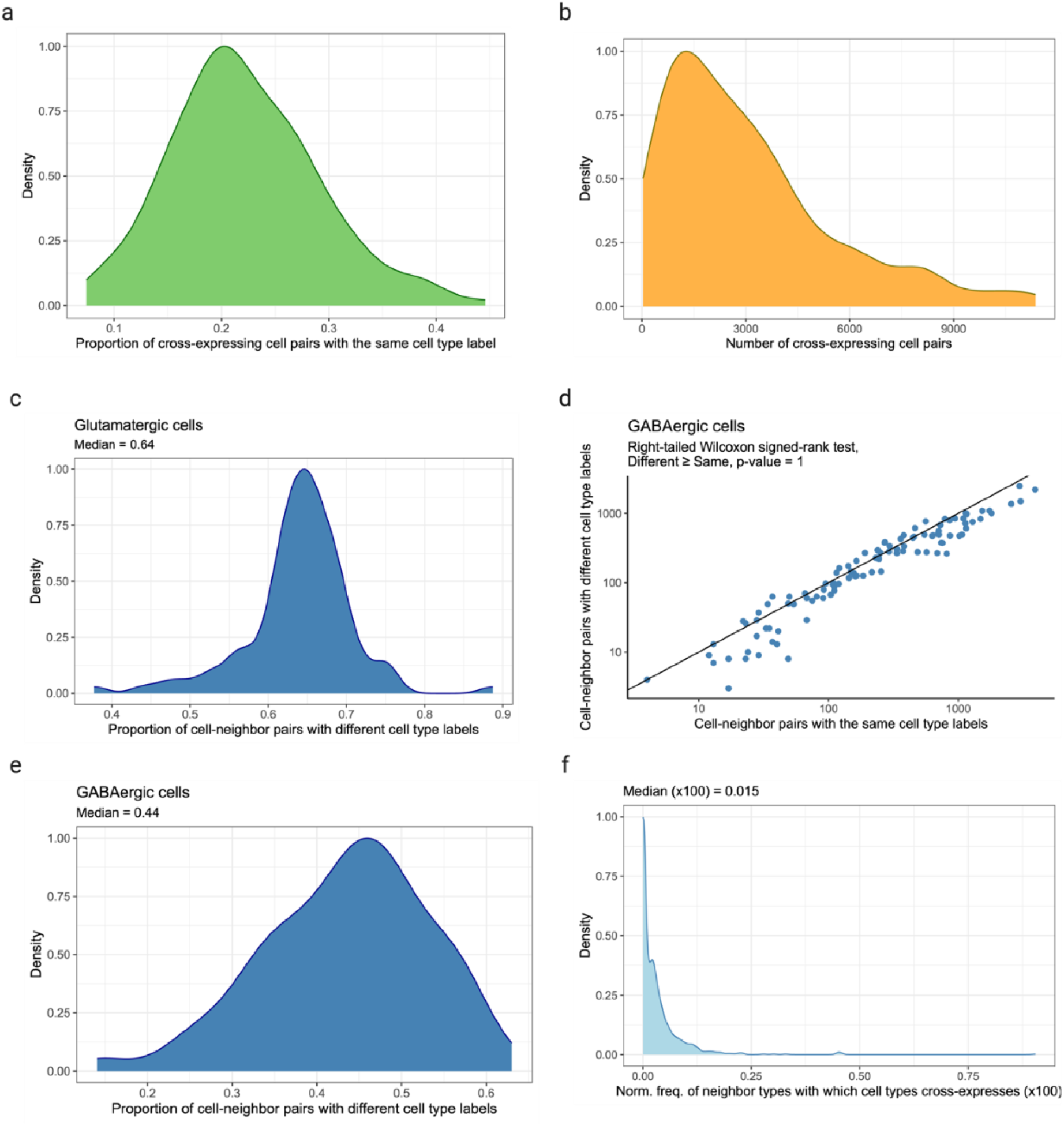
Relationship between cross-expression and cell type heterogeneity. **a**, Proportion of cross-expressing cell pairs belonging to the same cell type label. **b**, Number of cell-neighbor pairs involved in cross-expression. **c**, Proportion of cell-neighbor pairs with different cell subtype labels given that both were labeled ‘glutamatergic’ at the higher level in the cell type hierarchy. **d**, Number of cell-neighbor pairs with the same or different cell subtype label given that both were labeled ‘GABAergic’ at the higher level in the cell type hierarchy. Each point is a cross-expressing gene pair. **e**, Same as in (c) but for ‘GABAergic’ cells. **f**, Proportion of neighbor cell types against which cell types cross-express. Created with BioRender.com.

**Extended Data Figure 3:**
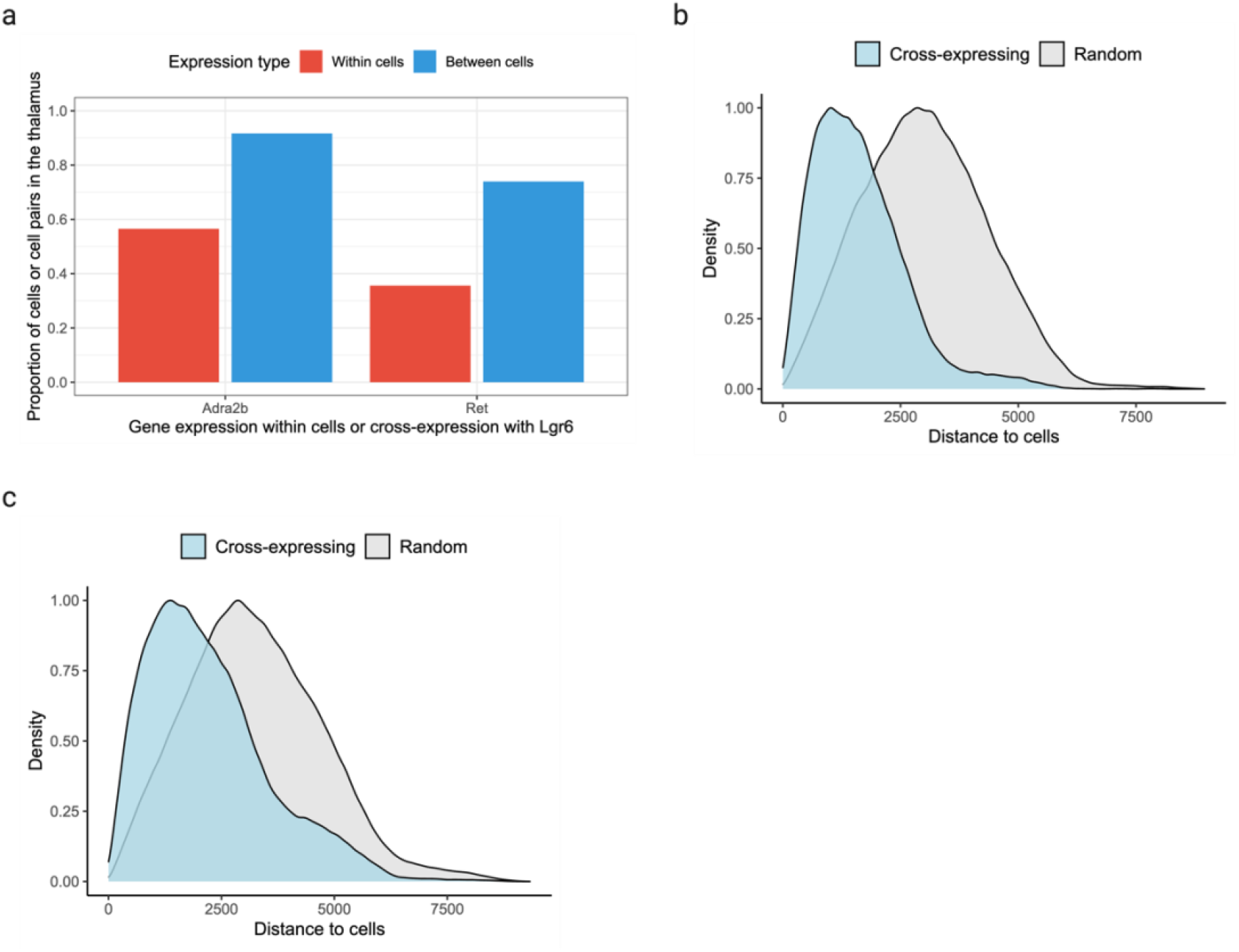
Combinatorial anatomical markers discovered using spatially enriched cross-expression. **a**, Proportion of *Adra2b*- and *Ret*-expressing cells in the thalamus (red) and the proportion of cell pairs in the thalamus (blue) when cross-expressing with *Lgr6*. **b-c**, Distances between cross-expressing cells versus those between cross-expressing and randomly chosen cells for genes *Lgr6* and *Adra2b* (c) and for *Lgr6* and *Ret* (d). Smaller distances mean that cross-expressing cells are nearer each other (spatial enrichment) than expected by chance (p-values ≤ 0.01, left-tailed Mann-Whitney U test). Created with BioRender.com.

**Extended Data Figure 4:**
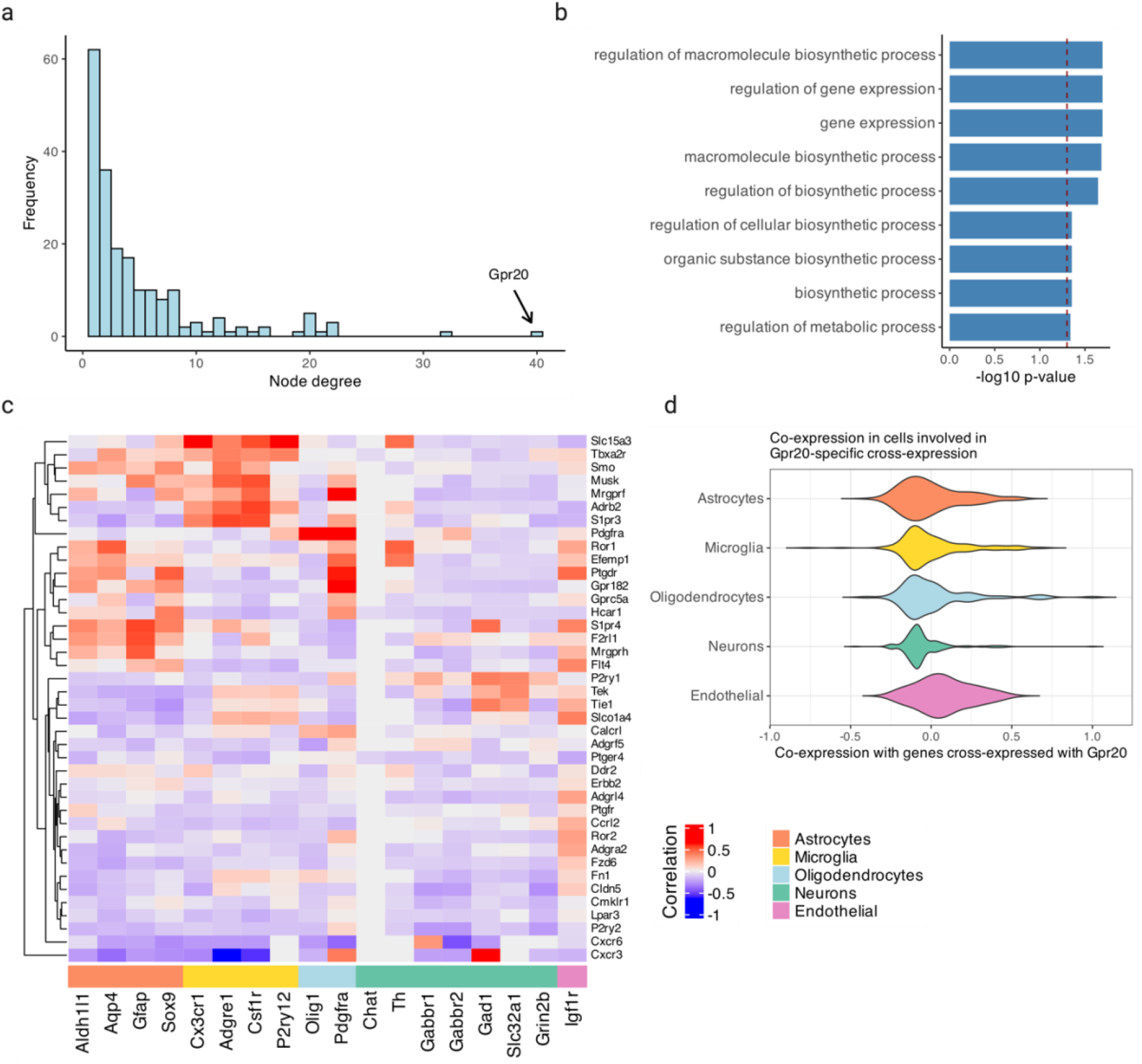
Exploration of *Gpr20* and its cross-expressing genes. **a**, Distribution of node degree, with *Gpr20* highlighted. **b**, Gene ontology (GO) functional groups for genes cross-expressed with *Gpr20*. **c**, Co-expression of genes cross-expressed with *Gpr20* (right) against cell type marker genes (bottom). For each gene, co-expression was computed using cells involved in cross-expression and not the entire dataset. **d**, Distribution of cell type marker genes’ co-expression across the genes in (c). Created with BioRender.com.

**Extended Data Figure 5:**
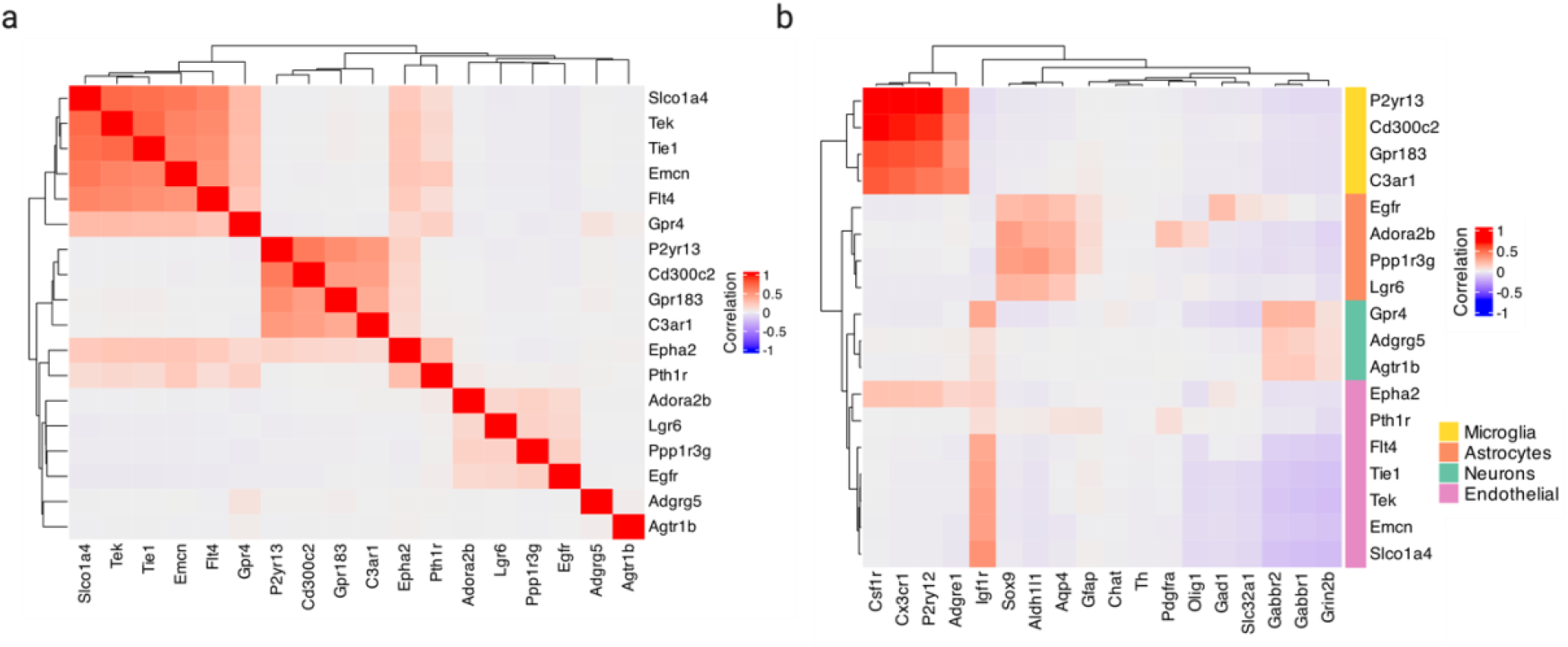
Exploration of the MERFISH cross-expression (sub)network. **a**, Co-expression of genes in the subnetwork. **b**, Co-expression between genes in the subnetwork (right) and cell type marker genes (bottom). Created with BioRender.com.

**Extended Data Figure 6:**
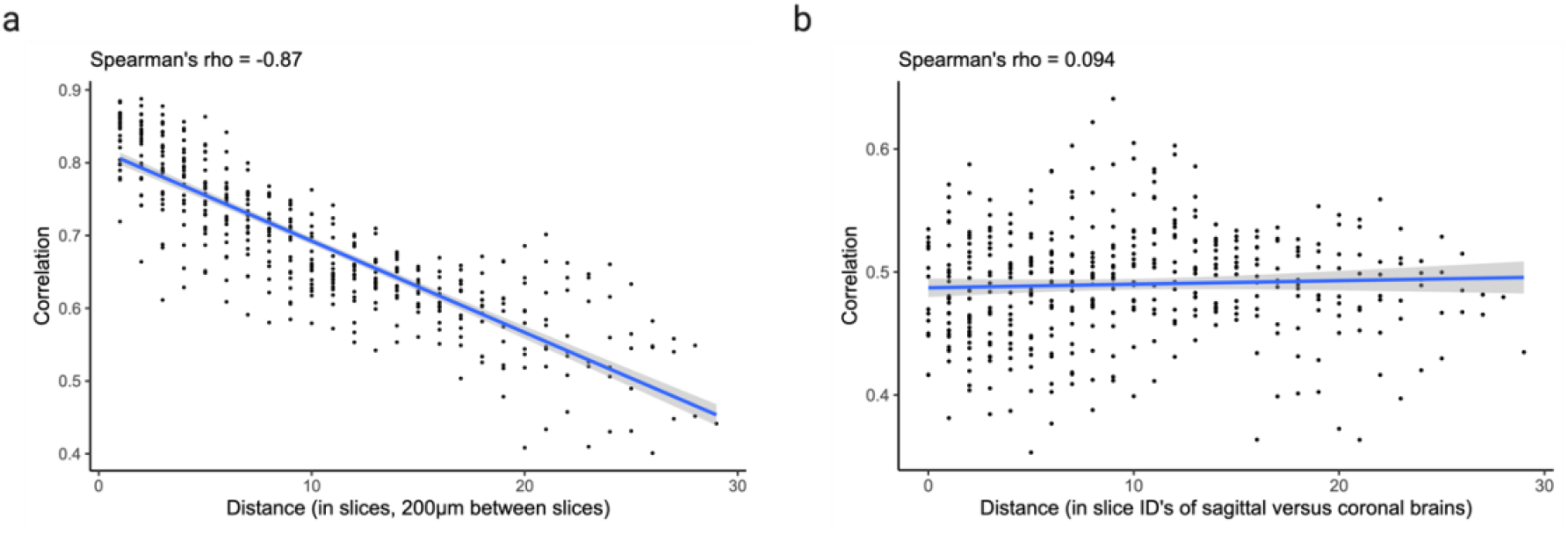
Cross-expression network similarity between slices. **a**, Slice-specific cross-expression networks compared and shown as a function of distance between slices. **b**, Same as in (a) but slice-specific networks compared between sagittal and coronal datasets, where the “distance” is the difference in slice ID’s. Shaded areas are 95% confidence intervals. Created with BioRender.com.

**Extended Data Figure 7:**
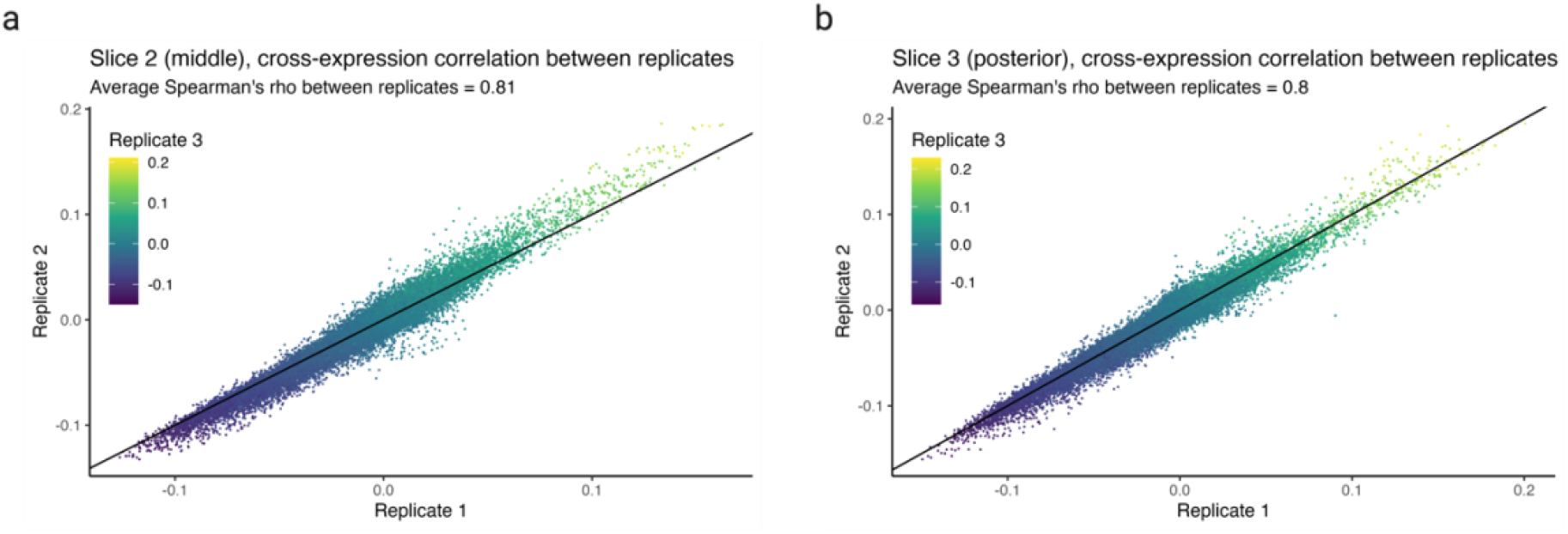
Cross-expression network similarity between replicates. **a-b**, Cross-expression networks compared between three replicates for the middle (a) and posterior (b) slices. Created with BioRender.com.

**Extended Data Figure 8:**
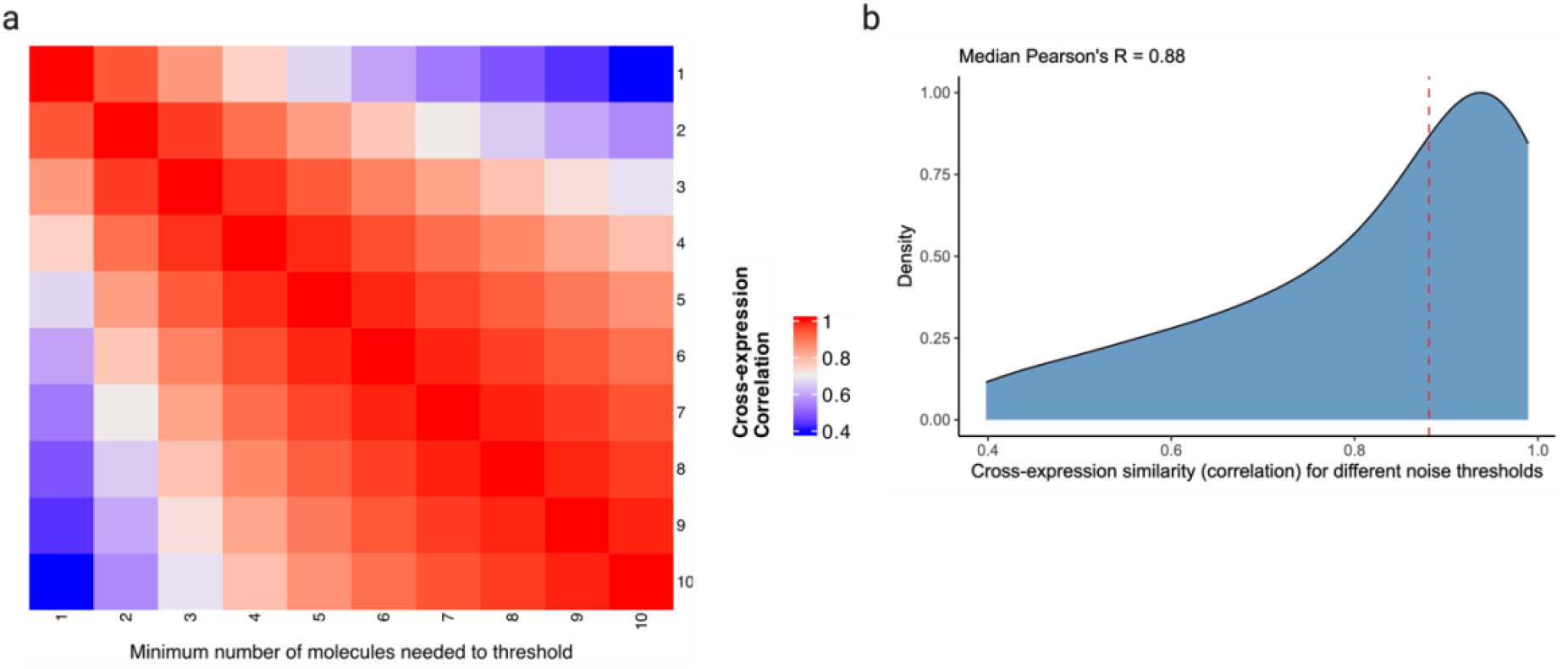
Cross-expression network similarity at different levels of gene expression noise thresholds. **a**, Cross-expression networks compared after applying different noise thresholds, which are the minimum number of molecules a gene must express within a cell to be considered as detected. **b**, Distribution of the network similarities across noise levels, with the median indicated using the dotted line. Created with BioRender.com.

**Extended Data Figure 9:**
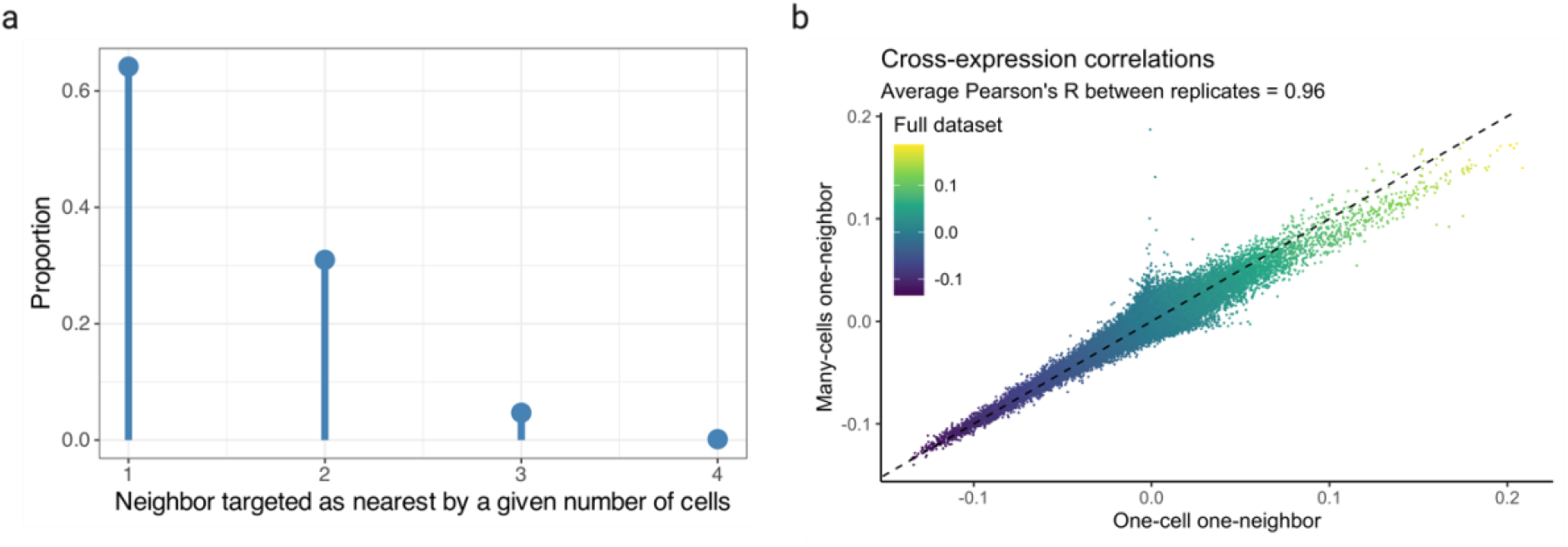
Patterns of cell-neighbor mappings and their relationship with cross-expression. **a**, Cells considered as ‘nearest neighbor’ by other cells reported as a proportion of total cell-neighbor relations. ‘1’ is one-to-one mapping and ‘2-4’ is many-to-one mapping. **b**, Cross-expression networks computed using one-to-one mappings, many-to-one mappings, and the full dataset (both mappings). Created with BioRender.com.

## Online Methods

We first explain the theoretical underpinnings of our approach and outline the features of the associated R package. We then specify how these are used in various analyses.

### Statistics of cross-expression between a gene pair

Cross-expression is the mutually exclusive expression of a gene pair across neighboring cells. To assess whether gene A’s expression in cells and gene B’s expression in their spatial neighbors is significant, we calculate the probability using the hypergeometric approach

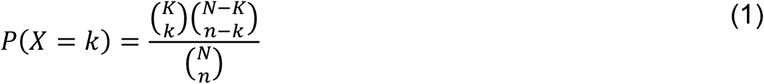

where *N* is the population size, *K* is the number of possible successes, *n* is the number of samples or draws, *k* is the number of observed successes, and the form 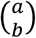 is the binomial coefficient.

Equation 1 outlines all the ways in which success can be observed—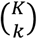—*and* (product rule) all the ways in which failure can be obtained—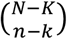—normalized by all possible ways of generating our sample 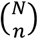, making the outcome probabilistic by bounding it between [0,1]. Traditionally, the *n* samples are assessed for the presence of some property *k*. Here, we sample cell-neighbor *pairs* and ask whether the cell expresses gene A while the neighbor expresses gene B. Thus, the sample size *n* is the number of pairs where the cells express gene A, the number of observed successes *k* is the number of pairs where the cells express gene A and the neighbors express gene B, and the number of success states *K* is the number of pairs where the neighbors express gene B. The population size *N* is the total number of cell-neighbor pairs, including those that co-express A and B and those that express neither gene. To calculate the probability of *k* or more successes, we use the hypergeometric cumulative distribution function (CDF)

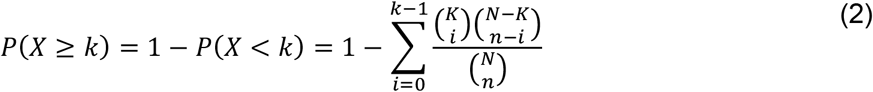

A value lower than alpha *α* indicates an unusually large number of pairs where the neighbors express gene B and the cells express gene A, making their cross-expression significant.

### Statistics of cross-expression between all gene pairs

We need to assess cross-expression across all gene pairs, which rise quadratically by 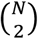 or 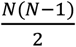 for *N* genes. To efficiently explore this space, we implement the procedure above using matrix operations and specialized packages in R.

We begin with a cells-by-genes expression matrix **E** and a cells-by-coordinates location matrix **L**, where the coordinates in our data are cell centroids on two-dimensional slices. We input **L** into `RANN` package’s function `nn2` with search type as `standard`, which implements a kd-tree algorithm to explore data subspaces and efficiently finds the *n*-th neighbors [89,90]. Using the neighbor indices, we re-order the expression matrix **E** to generate the neighbors-by-genes matrix **E′**. The value of *n* can be changed to generate paired gene expression matrices, where the corresponding rows represent cells and their *n*-th nearest neighbors. (To use a distance-based approach, one can use the *n*-th neighbor insofar as the average distance to this neighbor is smaller than the threshold.)

Our aim is to use **E** and **E′** to compute *N* (population), *K* (neighbors with B), *n* (cells with A), and *k* (neighbors with B when their corresponding cells express gene A) for each gene pair. These four values can be inputted into R’s `phyper` function for all gene pairs to calculate probabilities. The population size *N* is the total number of cells and is the same across all pairs. To compute *n*, we binarize **E** based on if the genes are expressed, and compute co-occurrences using the dot product

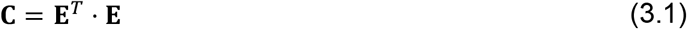

where **C**_*ii*_ is the number of cells expressing gene *i* and **C**_*ij*_ is the number of cells co-expressing genes *i* and *j*. We perform

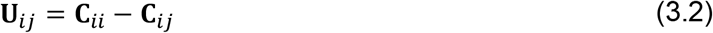

where **U**_*ij*_ is the number of cells uniquely expressing gene *i*. We implement this by extracting the diagonal of **C**, and “broadcast” it against its off-diagonal entries, thus aligning the corresponding values before subtraction. For each pair, this gives us the number of cells *n* uniquely expressing each gene. We perform an analogous calculation for *K* using **E′** instead of **E**, giving us the number of neighbors uniquely expressing each gene within a gene pair.

We now turn to *k*, the number of neighbors observed with gene B given that their corresponding cells express gene A. Using binarized matrices **E** and **E′**, we compute the number of cell-neighbor pairs such that the cells express gene A without gene B and the neighbors express gene B without gene A

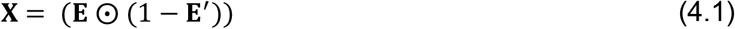

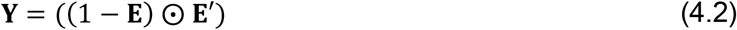

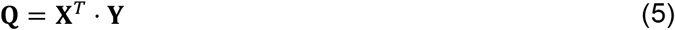

where ⨀ is the Hadamard (elementwise) product and **Q**_*ij*_ is the number of cell-neighbor pairs with mutually exclusive expression. In **X, E** contains ‘1’ in cells where a gene is expressed and 1 − **E**′ contains ‘1’ in neighbors where a gene is *not* expressed. Their elementwise product **X** has ‘1’ to indicate genes’ presence in cells and their absence in neighbors. **Y** shows the analogous procedure for genes’ presence in the neighbor and their absence in cells. Hence, the dot product of **X** and **Y** gives **Q**, a genes-by-genes asymmetric matrix, whose entries show the number of cell-neighbor pairs with mutually exclusive expression. (**Q** is asymmetric because the number of cell-neighbor pairs in the A-to-B and B-to-A directions are not always identical.) This is *k* or observed successes. These steps generate four number – *N, K, n*, and *k* – per gene pair. We input these into R’s `phyper` function in accordance with equation (2), giving us corresponding p-values.

Since **Q** is asymmetric, we obtain two p-values per gene pair, one in the A-to-B and the other in the B-to-A direction. We perform Benjamini-Hochberg false discovery rate (FDR) correction on the entire p-value distribution [91]. For each gene pair, we then use the lower FDR-corrected p-value as the final output **P** to indicate whether or not these genes cross-express.

### Cross-expression networks

We can threshold and binarize **P** at a pre-selected alpha *α* to form an adjacency matrix **N**, where ‘1’ indicates cross-expression (edges) between genes (nodes)

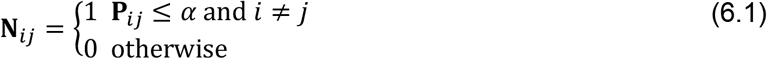

This allows us to perform cross-expression network analysis, where higher-order community structure is discovered using shared connections between genes

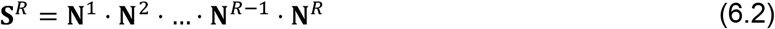

where we restrict *R* = 2 to discover second-order connections between genes.

### Cross-expression correlations

To compute correlations between genes, we first use the “masks” **X** and **Y** (eq. 4.1 and 4.2) and perform elementwise multiplication with non-binarized **E** and **E**′, respectively. This makes the gene counts “0” if cell-neighbor pairs co-express the gene pair of interest. We then compute the correlation between all pairs of columns (genes) of **E** and **E**′, yielding a gene-by-gene correlation matrix. We report the results as the average of this matrix with its transpose, giving a symmetric gene-gene correlation matrix as the output.

### Cross-expression at multiple length scales

Cross-expression is coordinated gene expression between neighboring cells. Yet, these patterns may be present at larger length scales, requiring us to understand associations between regions. To facilitate this, we smooth the expression of each gene in a cell by averaging it with its expression in *n* nearby cells. Using the `RANN` package [89,90], we find the indices of each cell’s *n* nearest neighbors, and make the corresponding values ‘1’ in the cells-by-cells matrix **C**

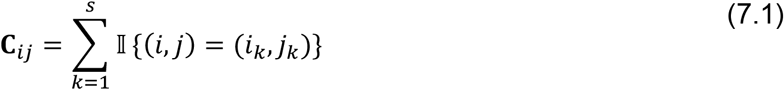

where the indicator function 𝕀 specifies

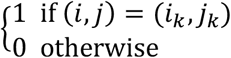

and

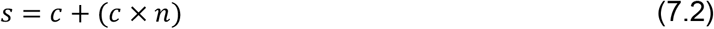

where *c* is the number of cells and *n* is the number of neighbors. Here, *s* is the total number of row-column indices *i*-*j* that *k* iterates over. We perform averaging using the expression matrix **E**

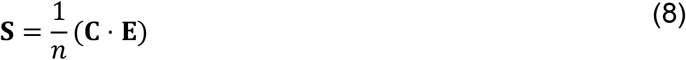

where **S**_*ij*_ is the *j*-th gene’s average value in *i*-th cell across *n* neighbors. The smoothed gene expression matrix **S** can be used for downstream analysis.

### Bullseye scores as effect size

The bullseye scores quantify the effect size by comparing cross-expression with co-expression. Here, the number of neighbors with gene B is compared to the number of cells co-expressing genes A and B. We use binarized cell and neighbor expression matrices **E** and **E′**, respectively

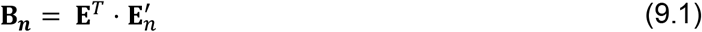

where *n* is the *n*-th neighbor, giving us *n* gene-by-gene asymmetric matrices **B**_***n***_. The *i*-th and *j*-th entries of **B**_***n***_ indicate the number of *n*-th nearest neighbors expressing gene B when cells in **E** express gene A. **B**_***n***_ is a co-occurrence matrix when *n* = 0. Viewing **B**_***n***_ as a tensor with dimensions *i, j*, and *n*, for each gene pair we take the cumulative sum and normalize across the neighbors

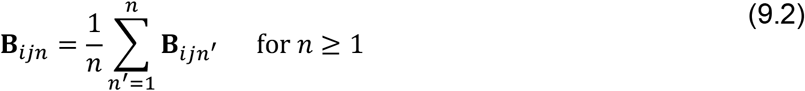

These matrices can be compared with **B**_*n*=0_ to find the ratio of cross-expression to co-expression and/or log_2_-transformed for further analysis. The output is provided as an array of matrices (tensor) or as an edge list, where columns represent different *n* neighbors.

### Expression of gene pairs on tissue

A powerful way of viewing cross-expression is to plot the cells and color them by their gene expression. For a gene pair, a cell can express genes A, B, both, or neither. We make these plots for user-selected gene pairs using the expression matrix **E** and the cell coordinates matrix **L**. We can also exclusively highlight cross-expressing cell-neighbor pairs. Finally, the tissue sections are often not upright, partly due to their misorientation with respect to the glass slide, making it difficult to interpret the results. Accordingly, we rotate them using user-defined *n*-degrees

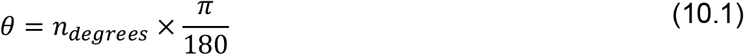

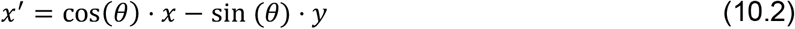

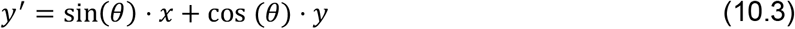

where *x*′ and *y*′ are the cell coordinates after counterclockwise rotation. Rotation does not change the distances between cells, so *x*′ and *y*′ can be used for downstream analysis.

### Spatial enrichment of cross-expression

Cross-expressing cells may be distributed across the tissue or show spatial localization. To quantify their enrichment, we first average the distance between cell-neighbor pairs. We next compare the distances between all cross-expressing cells to the distances between cross-expressing and randomly selected cells. If the former distance is significantly smaller than the latter distance, then cross-expression is spatially enriched.

### Data acquisition and preprocessing

#### BARseq datasets

We collected a sagittal mouse brain data (P56 male) from the left hemisphere (20µm thick sections, 300µm distance between slices) and measured 133 genes, including ligand-receptor pairs (neuropeptides, neuropeptide receptors, monoamine receptors such as cholinergic, adrenergic, serotonergic, and dopaminergic). This dataset includes 16 slices, which were assayed as previously described [35]. We removed cells expressing fewer than 5 genes or with less than 20 counts.

We downloaded the BARseq coronal mouse brain dataset [35], which contains 9 brains, with 4 control and binocular enucleation pairs as well as a pilot brain. These data contain 289 coronal slices and 109 genes. They were quality controlled just like the sagittal dataset.

#### 10× Genomics Xenium datasets

We obtained three datasets from the Xenium platform https://www.10xgenomics.com/datasets. The first dataset contains 1 coronal slice with 5006 genes obtained from https://www.10xgenomics.com/datasets/xenium-prime-fresh-frozen-mouse-brain. We removed cells with fewer than 50 counts.

The second dataset contains 3 coronal age-matched AD-control FFPE brain pairs, where the ages are 2.5 months, 5.7 months, and 13.4 (control) versus 17.9 months (AD). This data includes 346 genes and was obtained from https://www.10xgenomics.com/datasets/xenium-in-situ-analysis-of-alzheimers-disease-mouse-model-brain-coronal-sections-from-one-hemisphere-over-a-time-course-1-standard. We removed cells with fewer than 50 counts.

The third dataset includes 4 coronal replicates with 248 genes. Three of these were obtained from https://www.10xgenomics.com/datasets/fresh-frozen-mouse-brain-replicates-1-standard and the remaining from https://www.10xgenomics.com/datasets/fresh-frozen-mouse-brain-for-xenium-explorer-demo-1-standard. We removed cells with fewer than 30 counts.

#### EEL FISH dataset

We obtained a sagittal brain section data assaying 440 genes [92]. We removed cells with fewer than 15 counts.

#### NanoString CosMx dataset

We obtained a coronal brain section data from https://nanostring.com/products/cosmx-spatial-molecular-imager/ffpe-dataset/cosmx-smi-mouse-brain-ffpe-dataset/. This dataset assays 950 genes. We removed cells with fewer than 100 counts.

#### Resolve Biosciences Molecular Cartography dataset

We downloaded one hemi-coronal slice data from https://resolvebiosciences.com/open-dataset/?dataset=mouse-brain-2021, which assayed 99 genes. We removed cells with fewer than 30 counts.

#### Vizgen MERFISH brain receptor map datasets

We obtained Vizgen MERSCOPE’s mouse brain receptor map from https://info.vizgen.com/mouse-brain-data. This data contains three coronal slices from three replicates, with the middle slice covering the center of the brain. It assays 483 genes. We filtered cells with fewer than 50 counts and those lacking brain region annotations (see below). We registered the slices to the Allen CCFv3 (Common Coordinate Framework version 3) brain region atlas [61]. To facilitate this, we annotated the cells using Seurat [93]. Here, we created a Seurat object and used `SCTransform` with the `clip.range` between −10 and 10. We then ran Principal Component Analysis (PCA), setting the number of components to 30 and specifying the features as genes. Next, we used `FindNeighbors` and `FindClusters` with the resolution set to 0.3. The clusters are cell type labels, which help us identify brain structures during registration. For registration, we used QuickNii (v3 2017) [94] to linearly align the slice to the Allen CCFv3 atlas using discernible regions like the hippocampus and the ventricles as anchors. We then used VisuAlign (v0.9) [94] to non-linearly transform the slice to improve alignment with the atlas. This procedure assigns a brain region annotation to every cell.

#### STARmap dataset

The STARmap dataset contains 20 coronal, sagittal, or spinal sections, where 1022 genes were assayed [34]. We removed cells with fewer than 30 counts.

#### Allen Institute MERSCOPE atlas dataset

We obtained MERSCOPE dataset from the Allen Institute brain atlas, which contains 500 genes measured across 59 coronal slices [32]. We removed cells with fewer than 15 unique genes or lower than 40 counts.

#### MERFISH brain atlas datasets

The first dataset assays 1122 genes across 239 coronal and sagittal slices [33]. We removed cells in the top and bottom 1% by total counts.

The second dataset assays 374 genes across age-matched (4 weeks, 24 weeks, 90 weeks) controls and LPS-infected coronal brains, with 49 total slices [95]. (LPS is lipopolysaccharide bacterial infection.) We removed cells expressing fewer than 5 genes or with less than 20 counts.

#### Single-cell RNA-seq (scRNA-seq) and single-nucleus RNA-seq (snRNA-seq) datasets

We obtained the scRNA-seq [32] and snRNA-seq [77] data from the Brain Initiative Cell Census Network (BICAN) cell type atlases. These data were collected from dissected tissue regions, giving us the cells’ coarse anatomical origin. We removed cells with a doublet score of 30 or above and randomly selected 10,000 cells from each region for subsequent analysis.

### Ligand-receptor cross-expression

We aimed to find cross-expression between known ligand-receptor pairs. In our sagittal data, we selected two slices and within each slice we chose a cortical region. These choices were made randomly. In practice, we chose the visceral area (VISC) in slice 3 and the somatosensory nose region (SSp-n) in slice 5. Next, we selected the well-known neuropeptide somatostatin *Sst* and its cognate receptor *Sstr2* as the candidate pair. Finding their cross-expression significant, we show their expression on tissue and highlight cross-expressing cells. We also compute their bullseye scores and report them as a ratio of cross-to co-expression across 10 neighbors.

### Cross-expression and cell type heterogeneity

We explore the relationship between cross-expression and cell type heterogeneity using the BARseq coronal data [35]. First, we use *Gfra1* and *Foxp2* to highlight cross-expressing cells and map different cell types to distinct shapes. Second, we count the number of cross-expressing cell-neighbor pairs for numerous genes. Since each cell has a cell type label, we compute cell type purity as the proportion of pairs with the same label. Third, we use the cell type hierarchy to assess if cell-neighbor pairs with the same H1 label have the same H3 label. We first find cross-expressing gene pairs using the entire dataset. Next, using cell pairs with the ‘glutamatergic’ H1 label, we compute the number of pairs with the same or different H3 labels. We perform a similar analysis using cells labelled as ‘GABAergic’ at the H1 level. Finally, we compute the frequencies with which cell type label combinations are associated between neighboring cells and normalize this by the expected frequencies of those cell type pairs in the population.

### Replicability of cross-expression

We evaluate the replicability of our data using the hypergeometric and correlation-based approaches, which provide independent ways of quantifying cross-expression. We predict the binary p-values (significant or not, alpha = 0.05) using correlations and report the performance as AUROC. For performance within datasets, we predict the labels of a held-out slice using the average correlation of the remaining slices using a leave-one-out cross-validation procedure, with the results reported as AUROC against the number of slices/ samples used. For performance between datasets, we predict the labels of slices in one study using correlations of slices from another study and report the performance as median AUROC between study pairs. This process requires using gene pairs shared between studies’ panels.

### Creation and annotation of meta-analytic cross-expression network

Our cross-expression meta-analytic network included connections (edges) between genes (nodes) if they were cross-expressed in two or more studies. We used the `igraph` package [96] to create a network and Leiden clustered it [97], with “modularity” as the objective function ran for 100 iterations. Since Leiden clustering is somewhat stochastic, we performed this process 100 times, giving us 100 distinct partitions/ community assignments per gene. To find consensus communities, we calculated the adjusted Rand index (ARI) [98], which tells us the degree to which two clustering results agree with each other, between all pairs of partitions. Next, we found the average ARI for each partition and choose the one with the highest ARI, which gives us the ‘consensus’ communities. Importantly, we observed that multiple partitions had the same highest average ARI, and upon further inspection, we discovered that these partitions were identical, giving credence to our final community assignments.

After assigning genes to communities, we performed gene ontology (GO) [99,100] enrichment per community using the union of the studies’ gene panels as the background. We merged significant GO terms together using Revigo [101], which gave us the biological processes (BP), cellular components (CC), and molecular functions (MF) of each community. We performed minor manual curation of these terms to generate our final GO annotations. The meta-analytic network was visualized using Cytoscape (v3.10.1) [102,103].

### Networks of cross-expression when studying *Gpr20*

Using the MERFISH data, we computed cross-expression p-values between all genes and binarized the matrix at *α* ≤ 0.05 to create an adjacency matrix. We calculate the node degree as the number of edges formed by each gene and create a network with second-order edges (shared connections) as outlined in equation 6.2. We set the threshold for second-order edges to 4, meaning that two genes are connected if they share at least 4 first-order edges, ensuring that the higher-order network is robust. Next, we use the `igraph` package [96] to perform Louvain clustering [104] (with default parameters) on the second-order network and thus assign genes to communities.

We visualize the network using Cytoscape (v3.10.1) [102,103], mapping node size to degree, color to node community, and edge color to edge type (first-order, second-order, or both). We use the “organic” layout and apply “remove overlaps” from the `yFiles` app [103] and tweak the network to further reduce overlaps. Finally, we use the `Legend Creator` app [103] to render a legend with node degree size and community assignment.

Because our network revealed *Gpr20* as topologically salient, we performed gene ontology (GO) [99,100] enrichment analysis on genes that cross-expressed with it (‘test set’). Here, we used the entire gene panel (except *Gpr20*) as the background set and used the hypergeometric test to determine if it significantly overlapped with the test set, giving us p-values for each GO functional group. We report FDR-corrected p-values. Additionally, for each gene cross-expressed with *Gpr20*, we used the cells involved in cross-expression, rather than the entire dataset, to compute co-expression with cell type marker genes and compared these global profiles between marker types.

Since the cells expressing *Gpr20* visually showed spatial autocorrelation, we assessed their neighbors as well as randomly chosen cells for the expression of *Gpr20*. We L1-normalized the counts for both groups, rendering them into probability distributions, and computed cumulative sums. To calculate the area under curve (AUC), we scaled the neighbor order between 0 and 1 and used the `trapz` function from R’s `pracma` package to calculate the AUC [105].

Within the main network, we introduce a further constraint that cross-expressing genes must lack significant co-expression. We curate the subnetwork by removing genes with node degree of 1 and assign cell type labels based on genes’ co-expression with marker genes. Like before, we perform GO enrichment using the subnetwork genes as the test set and the gene panel as the background set, and report FDR-corrected p-values.

To assess whether cross-expression networks are more similar between adjacent slices than between distant slices, we compute slice-specific cross-expression networks and calculate Spearman’s correlation between these networks. The correlations are plotted against distances between slices, where the “distance” is the difference in the slice order. As a control, we compute the Spearman’s correlations between slice-specific networks obtained from different brains and plot this against the “distance” between the slice ID’s.

### Discovering combinatorial anatomical marker genes

We observed that cross-expression discovers anatomical marker genes that delineate the thalamus. To quantitatively assess this, we made a mask by combining the following regions: anterior group of the dorsal thalamus (ATN), intralaminar nuclei of the dorsal thalamus (ILM), lateral group of the dorsal thalamus (LAT), medial group of the dorsal thalamus (MED), midline group of the dorsal thalamus (MTN), ventral group of the dorsal thalamus (VENT), and ventral posterior complex of the thalamus (VP). Importantly, we compared every brain region annotation in our data with Allen CCFv3 atlas [61] and judged the ones presented here to best mark the thalamic regions. This allowed us to calculate the number of cells expressing each gene within or outside the thalamus. For cross-expressing cells, we considered a pair as thalamic if both cells were part of the regional mask. More generally, potential combinatorial markers can be discovered by assessing if their cross-expression is spatially enriched.

Our second exploration involved well-known genes *Foxp2* and *Cdh13*, which mark cortical layer 6 and show pan-layer expression in the cortex, respectively. These genes exhibited significant cross-expression, which was spatially enriched in layer 6, whose boundaries we identified using H2 cell type annotation. The spatial enrichment was viewed by comparing tissue plots with and without highlighting cross-expressing cells.

### Robustness of cross-expression to batch effects

To assess the replicability of the cross-expression signature, we used the MERFISH dataset containing 3 biological replicates (mouse brains) with 3 slices each, where the slices are sampled from approximately the same location across the brains. We compared the slice-specific networks between corresponding slices. Moreover, for slice-specific and brain region-specific networks, we performed comparisons within the BARseq sagittal data and within the coronal data [35] as well as between these two datasets. Finally, observing that the dorsal to ventral direction is sampled in both the coronal and the sagittal brains, we compared the densities of cross-expressing cells in the dorsal to ventral directions across these datasets.

### Cell segmentation quality control assessment

We assessed the quality of cell segmentation at a global level by comparing co-expression between the scRNA-seq [32] and MERFISH spatial transcriptomic data. Since the scRNA-seq was obtained from dissected brain regions, we established correspondence between these and the brain region annotations in the MERFISH data. The regions used in both data are reported in Supplementary Table 1. We included only those genes – and in the same order – as present in the MERFISH data. We calculated gene co-expression using Pearson’s correlation and compared these across the two datasets.

To quantify variability between platforms, we compared gene co-expression between scRNA-seq and snRNA-seq [77] for the same genes as above. Because the snRNA-seq was obtained from dissected brain regions, we established correspondence between these and the scRNA-seq data. The regions used in these data are reported in Supplementary Table 2. Like before, we quantified co-expression using Pearson’s correlation and compared it across the two datasets.

### Gene expression noise thresholds and cell-neighbor relations

Because gene expression measurement is noisy, we applied thresholds of 1 to 10 molecules, thus specifying the minimum number of counts per cell a gene must have to be considered expressed. We then compared cross-expression networks across these thresholds.

Additionally, a cell might be the nearest neighbor of one or more cells. To ensure that our framework captures this variability, we compare cross-expression networks for the one-to-one and many-to-one mappings with each other and with that of the full dataset.

### Benchmarking the algorithm’s speed

We assessed the speed of the cross-expression algorithm by duplicating our BARseq coronal data, where the gene panel ranged from 2,000 to 8,000 and the number of cells ranged from 20,000 to 200,000. We ran the cross-expression algorithm and calculated the time on a 16 GB Apple M1 Pro macOS Sonoma 14.5 laptop.

## Declarations

### Ethics approval and consent to participate

The experimental work in this study involved the collection of a mouse sagittal brain dataset using BARseq. All experimental procedures were carried out in accordance with the Institutional Animal Care and Use Committee at the Allen Institute for Brain Science.

### Consent for publication

Not applicable.

### Availability of data and materials

The datasets used in this study are available in the repositories or articles outlined in the “data acquisition and preprocessing” section. The BARseq sagittal data’s sequencing images are being deposited to Brain Image Library (BIL). While it is being approved, we stored the cell-level gene expression and cell metadata at https://drive.google.com/drive/folders/1fk5JDeVJcE71iH1AalCT0il9PN9DTJJm?usp=drive_link.

### Competing interests

L.F. owns shares in Quince Therapeutics and has received consulting fees from PeopleBio Co., GC Therapeutics Inc., Cortexyme Inc., and Keystone Bio. The remaining authors declare no competing interests.

### Funding

A.S. received funding from University of Toronto’s FAST Doctoral Fellowship, Ontario Graduate Scholarship, and NSERC’s Canada Graduate Scholarship. X.C. was supported by grants R01MH133181 and DP2MH132940. L.F. received support from R01MH133181 and J.G. from both R01MH133181 and R01MH113005. The content is solely the responsibility of the authors and does not necessarily represent the official views of the National Institutes of Health.

### Authors’ contributions

J.G. conceived the project. A.S. conducted the analyses, developed the software, and wrote the manuscript under supervision from J.G. M.R and H.C. collected the sagittal brain data under supervision from X.C. L.F. managed, curated, and parsed the datasets. S.C. created the R package. All authors interpreted the results and edited the manuscript.

## Acknowledgements

Not applicable.

## Authors’ information (optional)

Not applicable.

## Code availability

The R package is available at https://github.com/gillislab/CrossExpression/, including installation guide via Bioconductor and a tutorial with an example expression matrix and metadata.

## Notes

https://github.com/gillislab/CrossExpression

https://www.10xgenomics.com/datasets

https://info.vizgen.com/mouse-brain-data

https://nanostring.com/products/cosmx-spatial-molecular-imager/ffpe-dataset/cosmx-smi-mouse-brain-ffpe-dataset/

https://resolvebiosciences.com/open-dataset/?dataset=mouse-brain-2021

### Competing Interest Statement

Leon French owns shares in Quince Therapeutics and has received consulting fees from PeopleBio Co., GC Therapeutics Inc., Cortexyme Inc., and Keystone Bio. The remaining authors declare no competing interests.

### Summary of Updates

The abstract, introduction, results section 1, and large parts of the methods have been extensively rewritten. Two entirely new sections have been added in the results. The updated version now analyzes multiple datasets (13 total spanning ~25 million cells assayed using 8 technological platforms across 695 individual brain slices/ samples).

